# Lateral olivocochlear neurons modulate cochlear responses to noise exposure

**DOI:** 10.1101/2024.03.04.583418

**Authors:** Austen A. Sitko, Michelle M. Frank, Gabriel E. Romero, Lisa V. Goodrich

## Abstract

The sense of hearing originates in the cochlea, which detects sounds across dynamic sensory environments. Like other peripheral organs, the cochlea is subjected to environmental insults, including loud, damage-inducing sounds. In response to internal and external stimuli, the central nervous system directly modulates cochlear function through olivocochlear neurons (OCNs), which are located in the brainstem and innervate the cochlear sensory epithelium. One population of OCNs, the lateral olivocochlear (LOC) neurons, target spiral ganglion neurons (SGNs), the primary sensory neurons of the ear. LOCs alter their transmitter expression for days to weeks in response to noise exposure (NE), suggesting that they are well-positioned to tune SGN excitability over long time periods in response to auditory experience. To examine how LOCs affect auditory function after NE, we characterized the transcriptional profiles of OCNs and found that LOCs exhibit transient changes in gene expression after NE, including upregulation of multiple neuropeptide-encoding genes. Next, by generating intersectional mouse lines that selectively target LOCs, we chemogenetically ablated LOC neurons and assayed auditory responses at baseline and after NE. Compared to controls, mice lacking LOCs showed stronger NE-induced functional deficits one day later and had worse auditory function after a two-week recovery period. The number of remaining presynaptic puncta at the SGN synapse with inner hair cells did not differ between control and LOC-ablated animals, suggesting that the primary role of LOCs after NE is likely not one of protection, but one of compensation, ensuring that SGN function is enhanced during periods of need.

## INTRODUCTION

The sense of hearing originates in the cochlea, whose output depends on interactions among mechanosensitive hair cells, spiral ganglion neurons (SGNs), and efferent innervation from the central nervous system. As the primary sensory afferent neurons of the auditory system, SGNs directly transmit auditory information received from sensory inner hair cells (IHCs) to the central nervous system (**Fig. 1A**). However, this system is more than a simple relay and must also adjust its sensitivity to variable sound contexts and experiences, including noxious stimuli. In the periphery, this modulation is primarily coordinated by olivocochlear neurons (OCNs), which reside in the brainstem and project axons into the sensory epithelium of the cochlea (**Fig. 1A**) (1–3). OCNs consist of two major cell types: lateral olivocochlear neurons (LOCs) densely innervate Type I SGN peripheral fibers and medial olivocochlear neurons (MOCs) innervate outer hair cells (OHCs), with sparse collaterals onto SGN peripheral processes (**Fig. 1A**) (4–8). In response to sound, MOCs rapidly and dynamically tune OHC control over cochlear sensitivity (9–13). LOCs are believed to modulate SGN activity (3, 14, 15), but the precise nature of their synapses with SGNs and the functional consequences of this connection remain unclear.

**Figure 1.**
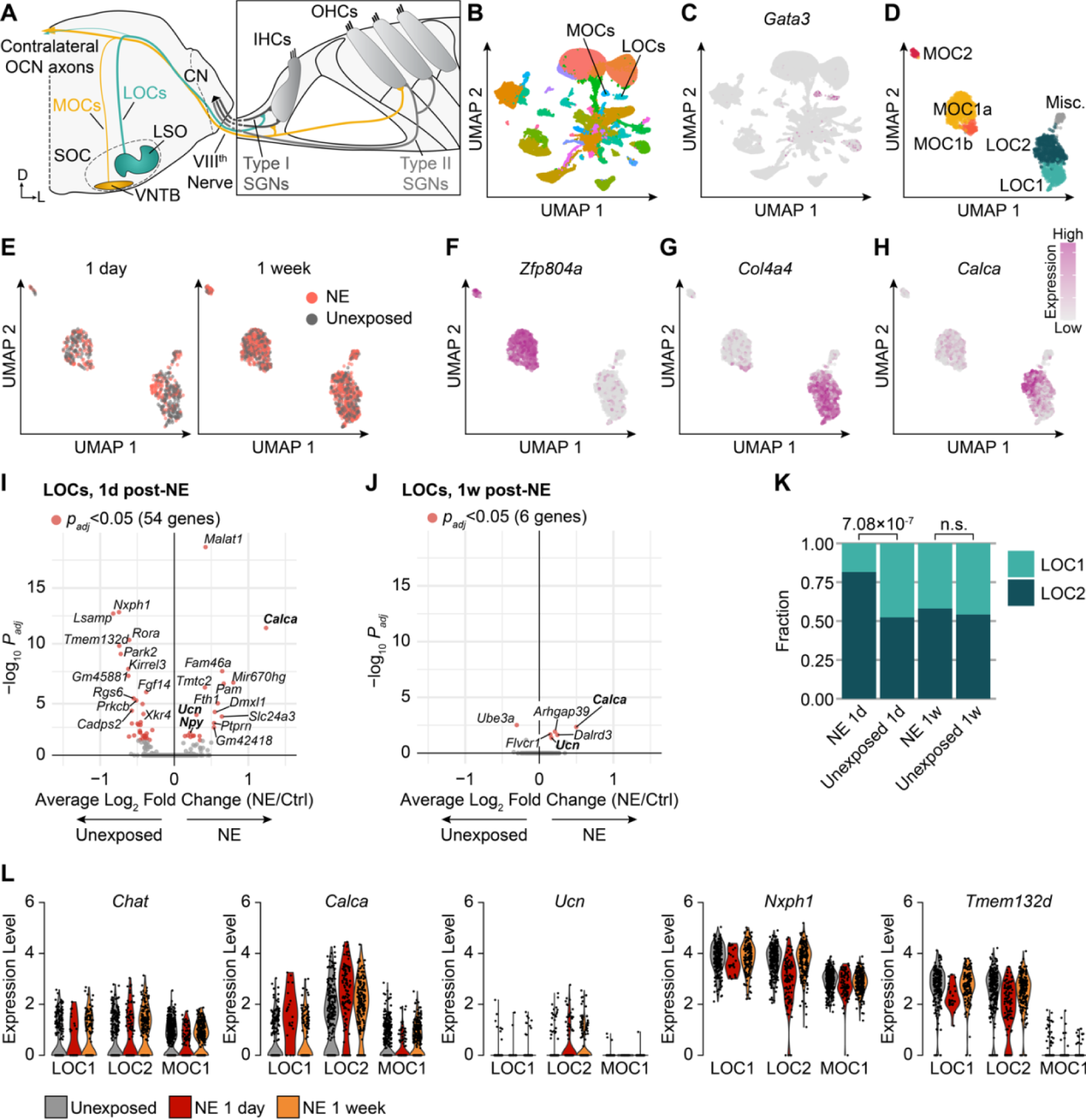
Single-nucleus RNA sequencing of olivocochlear neurons reveals activity-dependent transcriptional changes. **(A)** Schematic denoting major cell types and projections of the peripheral auditory system. Cell bodies of medial (MOC) and lateral (LOC) olivocochlear neurons (OCNs) reside in the superior olivary complex (SOC) within the ventral nucleus of the trapezoid body (VNTB) and the lateral superior olive (LSO), respectively. In the cochlea, sounds are transduced by inner (IHC) and outer (OHC) hair cells, which are contact by Type I and Type II spiral ganglion neurons (SGNs), respectively. SGN axons project into the cochlear nucleus (CN) in the brain. LOC axons synapse onto Type I SGNs, whereas MOCs primarily contact OHCs. D, Dorsal; L, Lateral. **(B, C)** UMAP representation of clustered single-nucleus sequencing dataset of 99,519 cholinergic brainstem neurons. OCNs are distinguished by their co-expression of motor neuron markers and the transcription factor *Gata3*. **(B)** Feature plot showing log-normalized expression of *Gata3*. For visualization purposes, expression scale is capped at the 95^th^ percentile. Expression scale is shown in **(H)**. **(D–H)** Sub-clustering results of 1,757 *Gata3*+ nuclei bioinformatically identified as OCNs. **(D)** UMAP representation of OCN subclusters, including two main categories of LOCs, three subclusters expressing MOC markers, and a miscellaneous cluster expressing markers for neither OCN type. **(E)** Distribution of LOCs with a history of noise exposure (NE) in the LOC subclusters was affected 1 day after NE, but clustering results overall were not driven by NE history. (**F–H)** Feature plots denoting log-normalized expression values of previously validated MOC **(F)** and LOC **(G)** markers. *Calca* **(I)** is expressed in both MOCs and LOCs but is highest in LOC2s. **(I, J)** Differentially expressed genes in LOCs 1 day (1d) **(I)** or 1 week (1w) **(J)** after NE. 54 genes were differentially expressed 1d after NE (Wilcoxon rank-sum test with Bonferroni *post-hoc* adjustment, *p_adj_* < 0.05), including peptides previously shown to be regulated by NE, labeled in bold. By 1w after NE, only 6 genes were differentially expressed and the magnitude of altered peptide expression is lower than at 1d post-NE. **(K)** The fraction of LOCs classified as LOC1 vs. LOC2 was significantly different 1d after NE (*p_adj_* = 7.07×10^-7^, Fisher exact test with Bonferroni *post-hoc* adjustment) but not 1w after NE (*p_adj_* > 0.999). **(L)** Violin plots denoting expression of indicated genes across subclusters and exposure conditions. Although *Chat* expression was unchanged across conditions, peptides such as *Calca* and *Ucn* increased in LOCs after exposure, whereas many membrane proteins, such as *Nxph1* and *Tmem132d* decreased in expression.

Both major OCN groups are implicated in protecting the auditory system from acoustic trauma. Excessively loud sounds can damage the cochlea, leading to temporary or permanent hearing loss depending on the duration and intensity of the stimulus. Cochlear supporting cells, hair cells, SGNs, or the synapses that link them can all be damaged—or completely eliminated— following traumatic noise exposure, resulting in elevated hearing thresholds, reflective of worse sound detection (16–21). If the noise exposure is sufficiently intense, both the loss of cochlear cells and the consequent increase in auditory thresholds will be permanent. After a more moderate noise exposure, much of this damage can be repaired, with thresholds returning to baseline after two weeks. Even after thresholds recover, however, there can be a permanent loss of synapses between IHCs and SGNs (22). As a result, fewer SGNs are activated by sound stimuli and thus the detection of auditory responses is diminished.

MOC-mediated feedback can reduce damage from traumatically loud sounds in a reflexive manner (13, 22–26), likely by decreasing OHC-mediated cochlear gain. LOCs, which generally act on slower timescales than MOCs, have also been implicated in protecting auditory function following an acoustic insult (25, 27, 28), presumably by decreasing SGN sensitivity. However, the role that LOCs play in acoustic trauma remains ambiguous, as prior studies were unable to definitively attribute observed effects specifically to LOCs due to the lack of tools to manipulate LOC signaling without affecting either some fraction of MOCs or neighboring, non-efferent auditory neurons in the brainstem.

Further complicating investigations into the functional role of LOCs, previous studies have observed opposing effects on SGN activity following indirect stimulation of LOCs (14) or application of neurotransmitters expressed by LOCs (29–31), indicating that these neurons likely have varied effects in the cochlea. Indeed, LOCs express a variety of neurotransmitters, neuromodulators, and neuropeptides, several of which are expressed heterogeneously (5, 32). Moreover, exposure to noise can induce changes in the expression profile of LOC signaling molecules that persist for days to weeks (5, 15, 33). These findings raise the possibility that LOCs influence cochlear function in multiple ways that can change in response to auditory experience.

To better understand the type of feedback that LOCs provide to the cochlea, we characterized their impact on hearing after exposure to excessively intense sounds. Single-nucleus RNA-sequencing (snRNA-seq) after noise exposure revealed that LOCs undergo transient changes in gene expression that are predicted to alter their signaling output for at least a week. Selective ablation of LOCs using a new intersectional chemogenetic approach revealed that LOCs suppress cochlear responses to very high-frequency sounds. Additionally, LOC-ablated animals had weaker auditory responses following noise exposure compared to mice with an intact LOC system, even though there was no effect on the number of pre-synaptic ribbons in IHCs. These findings suggest that LOCs enhance auditory function by modulating cochlear output in an experience-dependent manner.

## RESULTS

### Noise exposure induces transient transcriptional changes in LOCs

LOCs comprise a heterogeneous population that changes its molecular properties in response to traumatic levels of noise. A deeper understanding of the acute response to excessive noise stimuli can provide insight into how LOCs may function in a variety of other contexts. Therefore, we examined the transcriptional profiles of OCNs from 6–8-week-old (6–8w) mice exposed to a broadband 8–16 kHz stimulus at 110 dB sound pressure level (SPL) for 2 hours (2h), compared with unexposed littermates 1 day (1d; *N* = 9 exposed, 8 unexposed mice) and 7d (*N* = 15 exposed, 14 unexposed mice) following the exposure. This noise exposure (NE) stimulus has been found to upregulate TH, Ucn, NPY, and CGRP (5, 15), and similar exposure paradigms are associated with permanent deficits in auditory function (18, 34). We used *Chat^CreΔNeo^;Rosa26^Sun1-GFP^*mice to drive expression of a nuclear-localized GFP reporter in cholinergic neurons and dissected a region in the ventral brainstem that included OCNs and the nearby facial motor neurons. After dissociation, we enriched for GFP+ nuclei by fluorescence-activated cell sorting (FACS) and used the 10x Genomics platform to encapsulate and barcode single nuclei for sequencing.

After filtering out low-quality cells, our analysis is based on 99,519 total nuclei, including 1,757 cells bioinformatically identified as OCNs based on co-expression of *Gata3*, motor neuron markers like *Tbx20* and *Isl1*, and previously identified OCN markers like *Cadps2* (**Fig. 1B–C**). To further refine OCN cell types, we subclustered all nuclei identified as OCNs. Consistent with previous results (5), this analysis identified five main subclusters, including two main clusters that express MOC markers, two subtypes of LOCs, and a miscellaneous cluster that did not express either MOC or LOC markers (**Fig. 1D**). These subclusters also included an additional subtype of MOC that we had not identified previously, which we label here as MOC1b. Because this subcluster constitutes only a minority population of MOCs (53 MOC1b nuclei from unexposed animals versus 321 MOC1a) and there are relatively few differentially expressed genes between MOC1a and MOC1b clusters (**Fig. S1G**), we pooled these two MOC1 clusters for all subsequent analysis. In unexposed animals, markers for each of these subtypes are similar to those identified previously (5): the MOC2 subtype is distinguished from MOC1s by high levels of *Tox*, *Nkain2*, and *Nkain3* (**Fig. S1H**); the Misc. subtype is characterized by high expression of *Spef2* (**Fig. S1I**); and LOC2s are distinguished from LOC1s largely by enriched expression of transcripts encoding neuropeptides (**Fig. S1J**). Cluster identity was not driven by variability in cell health or read depth (**Fig. S1A–B**).

Compared to unexposed mice, LOCs from mice collected 1d after NE had significantly elevated expression of *Calca*, *Ucn*, and *Npy* transcripts (**Fig. 1I**; Wilcoxon rank-sum test with Bonferroni *post-hoc* adjustment), consistent with our previous finding that expression of CGRP, Ucn, and NPY proteins are elevated after NE (5). In addition, LOCs also appeared to downregulate several transcripts, including a high number of genes that encode cell-adhesion and transmembrane proteins (**Fig. 1I**). Overall, there was a significant shift in the fraction of LOCs classified as peptide-high LOC2s 1d after NE relative to unexposed controls (**Fig. 1K**). Given the limited number of bioinformatically identified LOC1s at 1d post-NE (24 LOC1s versus 106 LOC2s), we were unable to make reliable quantitative assessments about NE-dependent changes within each LOC subtype. Qualitatively, however, some genes—including *Calca*, *Nxph1*, and *Tmem132d*—appeared to undergo similar changes in both LOC1s and LOC2s (**Fig. 1L**). By 7d post-exposure, the majority of these gene-expression changes were no longer detectable (**Fig. 1J**). Although both *Calca* and *Ucn* transcripts remained significantly elevated 7d after NE, the magnitude of their expression change was lower than at 1d (**Fig. 1I, J, L**). Similarly, the relative fraction of neurons in the LOC2 cluster at 7d post-NE was comparable between NE and unexposed control mice (**Fig. 1K**), further suggesting that the transcriptional changes identified at 1d after NE are transient. Only minor differences were detectable between control animals collected at different times, indicating that these NE-dependent changes in LOC gene expression cannot be explained by batch effects or collection artifacts (**Fig. 1E, S1C**). In contrast, transcriptional changes in MOCs following NE were indistinguishable from batch-to-batch variation also present in control animals (**Fig. S1D–F**).

### Genetic strategy for specifically targeting LOCs

The finding that LOCs—but not MOCs—exhibited changes in expression of neural signaling molecules following NE (**Figs. 1E, S1D–F**), combined with prior findings of long-lasting changes in protein expression of such molecules (5, 15, 33), suggested a distinct role for LOCs in modulating cochlear properties following NE. To test how LOCs affect auditory function after NE, we developed an intersectional genetic approach to selectively access LOCs bilaterally, without impacting MOCs or other cells in the auditory brainstem.

Because LOCs express *Ucn* but MOCs do not (5, 35), we generated a *Ucn^Cre^* mouse using CRISPR/Cas9-driven homologous recombination to insert the *Cre* sequence in the endogenous locus of *Ucn* (**Fig. 2A**). During development, *Ucn* is initially expressed by LOCs throughout the lateral superior olive (LSO) before becoming restricted to LOC2 neurons in the medial wing in adults (5, 35). As a result, Cre is active in most LOCs in developing *Ucn^Cre^* mice. Indeed, LOC axons in the cochlea were robustly and selectively labeled when *Ucn^Cre^*drives recombination of *Rosa26^LSL-tdTomato^*, with no labeling of MOC axons or other cochlear cells (**Fig 2B–C**). Likewise, in the superior olivary complex, which houses the OCNs, most ChAT+ neurons in the LSO also expressed tdTomato, indicating efficient coverage of the LOC population (**Fig 2D–E**). By contrast, no MOCs were labeled. However, *Ucn* is also expressed in many other brain regions, so numerous tdTomato+ neurons were present throughout the brain (**Fig. 2F**). Thus, to further isolate LOC neurons, we leveraged the fact that the transcription factor *Gata3* is broadly expressed throughout much of the auditory system, including OCNs (36–38). Using a similar CRISPR/Cas-9 approach, we inserted the *FlpO* construct into the *Gata3* locus to generate *Gata3^FlpO^* mice (**Fig 2G**). As expected, there was widespread expression of tdTomato in the cochlea of *Gata3^FlpO^;Rosa26^FSF-tdTomato^*mice (**Fig. 2H**), as well as throughout the auditory brainstem and midbrain (**Fig. 2I–J**). Importantly, the nearby facial motor nucleus, which houses ChAT+, GATA3-motor neurons, was devoid of labeling in *Gata3^FlpO^;Rosa26^FSF-tdTomato^*mice (**Fig. 2J–K**).

**Figure 2.**
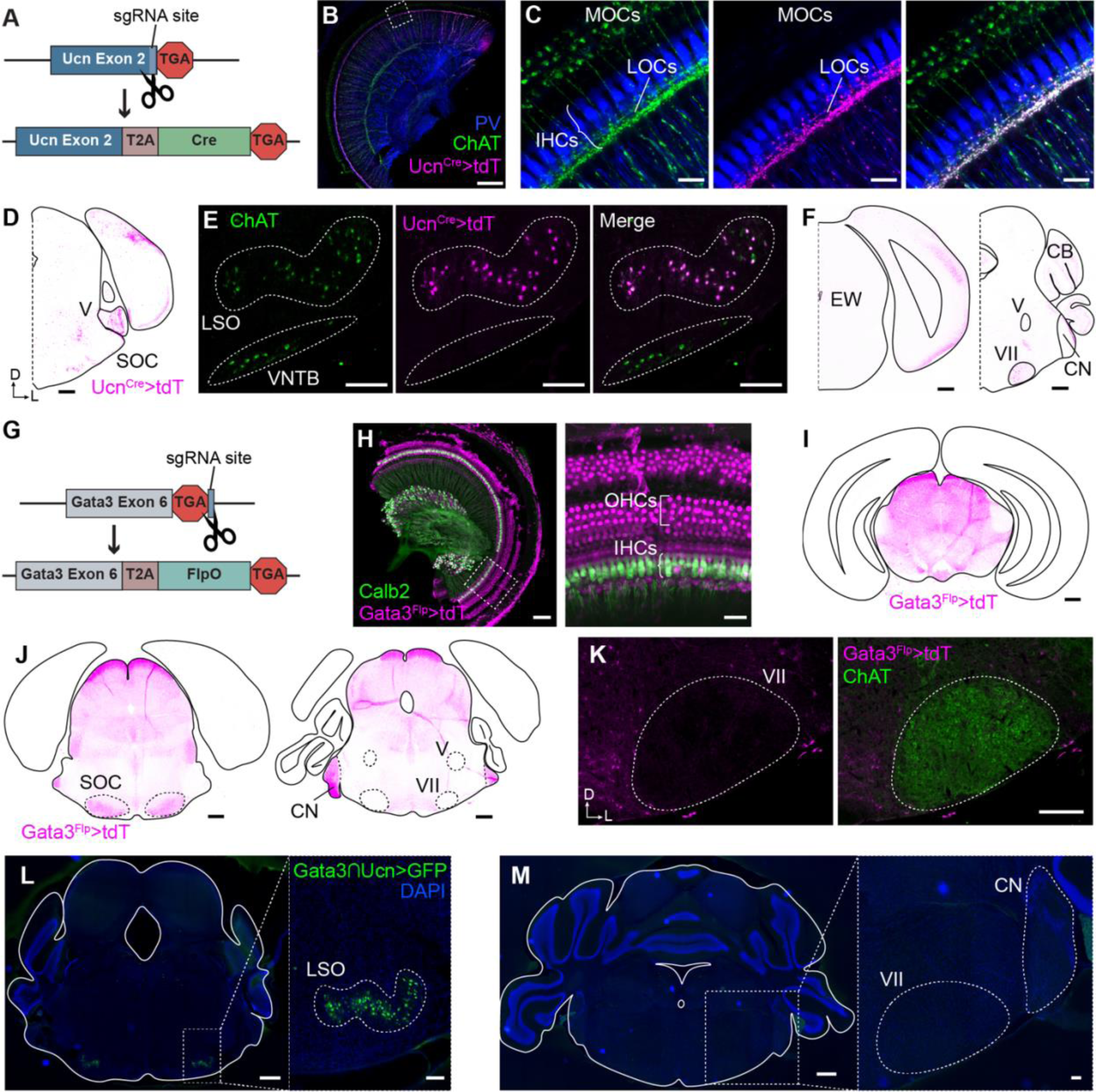
Generation of mouse lines to target LOCs. **(A)** Schematic of *Ucn^Cre^*allele. CRISPR/Cas9 was used to insert a T2A peptide and nuclear-localized *Cre* sequence immediately after the *Ucn* coding sequence, prior to the endogenous stop codon. **(B–F)** Expression of Ucn^Cre/+^; Rosa26^LSL-tdTomato/+^ (Ucn^Cre^>tdT, magenta) in the cochlea **(B, C)** and brain **(D–F)** of a P27 mouse. Anti-ChAT immunolabeling (green) defines both MOCs and LOCs. **(B, C)** Standard-deviation projections of confocal z-stacks showing tdTomato expression in the middle turn of the cochlea. TdT fluorescence colocalized with anti-ChAT immunolabel in the IHC region (labeled with Parvalbumin (PV), blue; curved brackets). No tdT signal was observed in MOCs projecting to the OHC region, indicating that *Ucn^Cre^*drives expression in LOCs but not MOCs. **(B)** Scale bar, 150 µm. **(C)** Higher-magnification image of designated region in **(B)**. Scale bar, 15 µm. **(D–F)** Coronal sections indicating Ucn^Cre^>tdT signal in various brain regions. Within the SOC, Ucn^Cre^>tdT signal was restricted to cholinergic cells of the LSO **(D, E)**. Ucn^Cre^>tdT signal was also visible in the trigeminal (V) and facial (VII) motor nuclei **(D, F)**; cortex **(D, F)**; Edinger-Westphal nucleus (EW, **F**); cerebellum (CB, **F**); and in fibers in the cochlear nucleus (CN, **F**). Scale bars, 500 µm **(D, F)**, 150 µm **(E)**. D, Dorsal; L, Lateral. **(G)** Schematic describing generation of *Gata3^Flp^* mouse line. **(H–K)** Expression of Gata3^Flp/+^; Rosa26^FSF-tdTomato/+^ (Gata3^Flp^>tdT, magenta) in the cochlea **(H)** and brain **(I–K)** of a P28 mouse. **(H)** Gata3^Flp^>tdT was found throughout the cochlea, including in Calb2+ spiral ganglion neurons (SGNs) and IHCs (green). Scale bars, 100 µm (left) or 25 µm (inset). **(I–K)** Gata3^Flp^>tdT was present throughout the midbrain and auditory brainstem, including the SOC and CN **(I, J).** No signal was detected in the cortex, cerebellum, or trigeminal and facial motor nuclei **(K).** Scale bars, 500 µm **(I, J)** or 250 µm **(K)**. **(L, M)** Intersectional reporter expression in 8-week-old *Ucn^Cre/+^; Gata3^Flp/+^; Rosa26^FLTG^* animals (*Gata3∩Ucn >GFP*, N = 3). GFP expression was found throughout the LSO **(L)** but is absent in the facial motor nucleus (VII) and CN **(M)**. Scale bars, 500 µm; inset, 100 µm.

Crossing the *Ucn^Cre^* and *Gata3^FlpO^*mice with the intersectional reporter *Rosa26^FLTG^*, in which GFP expression is induced in cells expressing both Cre and Flp recombinases (39), resulted in GFP expression in LOCs throughout the LSO, not just those that express Ucn as adults (**Fig 2L**). No GFP was expressed in nearby auditory regions, including the cochlear nuclei (CN) and other neurons in the superior olivary complex, as well as the facial motor nucleus (**Fig. 2M**). Sporadic GFP expression was detected in several other brain regions, with consistent signal in the midbrain and olfactory bulb (*N* = 3 8w mice). This intersectional genetic approach provides the first selective genetic access to LOCs, separately from MOCs, principal neurons of the LSO, and other cholinergic brainstem motor neurons.

### Intersectional chemogenetic ablation of LOC neurons

We next leveraged this genetic access to selectively ablate LOC neurons in adult mice and assess how loss of LOC input to the cochlea affected auditory function. To target LOC neurons for chemogenetic ablation, we crossed *Gata3^FlpO^;Ucn^Cre^;Rosa26^FLTG^*mice (*Gata3∩Ucn>FLTG*) with *Mapt^ds-DTR^* mice that express a dual-recombinase Flp*-* and Cre*-*dependent diphtheria toxin receptor (DTR). Cells expressing DTR are rendered capable of endocytosing diphtheria toxin (DTX), which in turn initiates cell death. Experimental mice that either expressed DTR in LOCs (i.e., *Gata3∩Ucn>FLTG,DTR* + DTX; LOC-ablated) or lacked DTR expression (i.e., *Gata3∩Ucn>FLTG* + DTX; DTX-control) received DTX. A second control group of animals carrying the DTR allele received saline (SAL) instead of DTX (i.e., *Gata3∩Ucn>FLTG,DTR* + SAL; SAL-control) (**Fig 3A**). Experimental and control mice received two intraperitoneal injections 72h apart of either DTX (50 μg/kg) or SAL (4 μL/g), depending on condition, between 4 and 7w of age (**Fig. 3B**).

**Figure 3.**
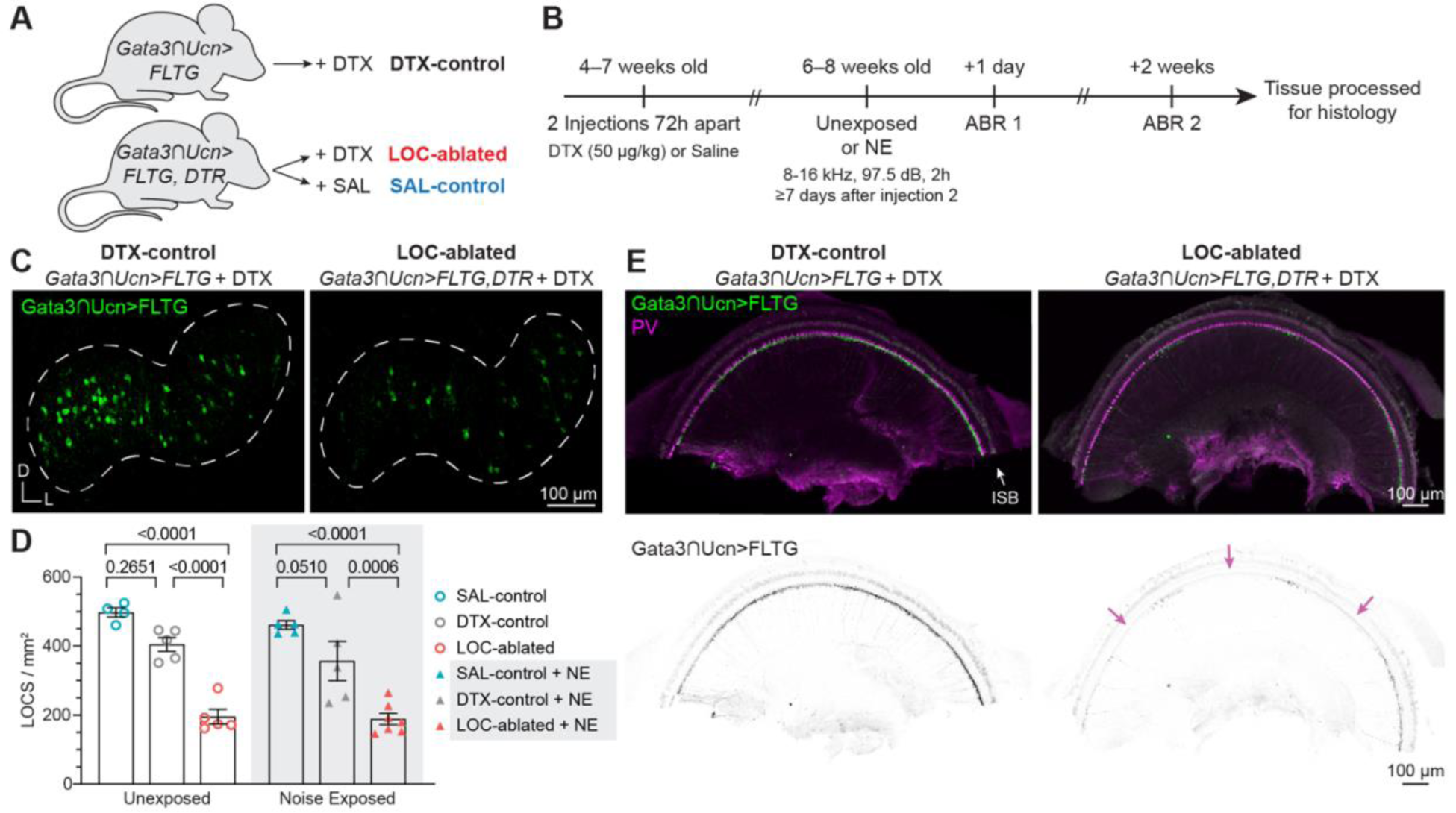
Intersectional chemogenetic approach to ablate LOCs. **(A)** Experimental conditions. In the DTX-control condition, *Gata3∩Ucn>FLTG* (*Gata3^Flp/+^;Ucn^Cre/+^;Rosa26^FLTG/+^;Mapt^+/+^*) mice received DTX. *Gata3∩Ucn>FLTG,DTR* (*Gata3^Flp/+^;Ucn^Cre/+^;Rosa26^FLTG/+^;Mapt^ds-DTR/+^*) mice received either DTX (LOC-ablated condition) or saline (SAL-control condition). **(B)** Experimental timeline. Male and female 4–7-week-old mice received two injections 72h apart of either DTX (50 μg/kg) or SAL (4 μl/g), depending on experimental condition. At 6–8 weeks old and ≥7 day following the second injection, mice were either noise exposed (NE) or left in home cages (Unexposed). Auditory function was assessed by ABR 1 day and 2 weeks later for all experimental groups. **(C–E)** Number of LOC somata in the LSO **(C)** and extent of LOC innervation in the cochlea **(D)** were reduced following chemogenetic ablation of LOCs. **(C)** LOCs marked by Gata3∩Ucn>FLTG GFP fluorescence (i.e., Gata3+,Ucn+ cells) in the LSO. There were fewer LOCs following ablation (right) compared with DTX-control (left) and SAL-control (data not shown) mice. See **Table 1** for mean ± SEM and sample sizes. **(D)** Quantification of LOC reduction. Unexposed and NE mice with ablated LOCs had significantly fewer LOC neurons (NE: 188.7 ± 16.40 LOCs/mm^2^, *N* = 7 mice; Unexposed: 195.4 ± 21.18 LOCs/mm^2^, *N* = 5) than either SAL-control (NE: 461.5 ± 12.44 LOCs/mm^2^, *N* = 5; Unexposed: 497.4 ± 13.62 LOCs/mm^2^, *N* = 4) or DTX-control mice (NE: 356.6 ± 56.97 LOCs/mm^2^, *N* = 5; Unexposed: 404.8 ± 19.76 LOCs/mm^2^, *N* = 5). Two-way ANOVA with Tukey’s multiple comparisons. Error bars, mean ± SEM. **(E)** Cochlea wholemounts (base turns) from DTX-control (left) and LOC-ablated (right) mice. SGNs and IHCs are labeled by PV (Parvalbumin) and LOCs axons are labeled with Gata3∩ Ucn>FLTG GFP fluorescence. LOC axon innervation of the cochlea was reduced in LOC-ablated mice (right, *N* = 17) compared to DTX-controls (left, *N* = 12), with large swathes of the ISB completely devoid of LOC axons (* in lower panels, inverted fluorescent signal). DTX = diphtheria toxin; SAL = saline; ABR = auditory brainstem response; NE = noise exposure; D = dorsal; L = lateral; PV = Parvalbumin; ISB = inner spiral bundle

**Table 1.** Number of LOCs/mm^2^.

| Condition | Exposure | LOCs/mm<sup>2</sup> | *n* (LSO sections) | *N* (mice) |
| --- | --- | --- | --- | --- |
| **SAL-Control**<br>*Gata3*∩*Ucn*>*FLTG*, *DTR* + SAL | Unexposed | 497.4 ± 13.62 | 23 | 4 |
|  | NE | 461.5 ± 12.44 | 30 | 5 |
| **DTX-Control**<br>*Gata3*∩*Ucn*>*FLTG* + DTX | Unexposed | 404.8 ± 19.76 | 30 | 5 |
|  | NE | 356.6 ± 56.97 | 40 | 5 |
| **LOCs-Ablated**<br>*Gata3*∩*Ucn*>*FLTG*, *DTR* + DTX | Unexposed | 195.4 ± 21.18 | 30 | 5 |
|  | NE | 188.7 ± 16.40 | 30 | 7 |

We validated the efficacy of this ablation approach histologically (**Fig. 3C–E**) and found that LOC-ablated mice had significantly fewer LOC neurons in the LSO (195.4 ± 21.18 LOCs/mm^2^, *N* = 5 mice, mean ± SEM) than either DTX-control (404.8 ± 19.76 LOCs/mm^2^, *N* = 5), or SAL-control mice (497.4 ± 13.62 LOCs/mm^2^, *N* = 4; **Fig. 3C–D**, **Table 1**). Qualitatively, there was no bias in the extent of LOC ablation across the medial–lateral or rostral–caudal extent of the LSO, suggesting a spatially uniform reduction in LOC neurons. Reduced LOC innervation was also apparent in the cochlea, where LOC axons beneath the IHCs were less dense and altogether absent in many areas (**Fig. 3E**). In sum, this intersectional chemogenetic approach allowed us to ablate a large fraction of LOCs selectively and bilaterally in the brainstem with a concomitant reduction of LOC input to the cochlea.

### Reduced LOC input to the cochlea enhanced responses to high-frequency stimuli

To determine how LOCs may fine-tune audition at the level of the cochlea, we first examined how baseline sound detection was affected in LOC-ablated animals. We assessed peripheral auditory function using the auditory brainstem response (ABR) test, which measures field potentials recorded near the inner ear in response to sound (see examples in **Figs. 4A & 5A**). ABR recordings provide a readout of neural activity as it propagates from the cochlea to brain regions along the auditory pathway. ABR thresholds indicate the sound intensity at which a neural response was first detectable along the ascending auditory pathway; the amplitude of wave I (location of wave I indicated by * in **Fig. 4A**, left panel) reflects the number of SGNs that responded synchronously to the sound stimulus. We found that DTX-control and LOC-ablated mice had similar baseline ABR thresholds (**Fig. 4A–B**), indicating that loss of LOC innervation did not affect baseline auditory sensitivity. Similarly, LOC-ablated and DTX-control mice had comparable wave I amplitudes at all frequencies except for the highest tested, 45 kHz. In response to 45 kHz tones, there was a significant leftward shift of the response curve in LOC-ablated mice compared to DTX-control animals, reflecting higher-amplitude responses when LOCs are depleted (**Fig. 4D**). Overall, these results indicate that decreased LOC input to the cochlea did not grossly alter normal auditory sensitivity, but increased either the recruitment or excitability of SGNs at 45 kHz.

**Figure 4.**
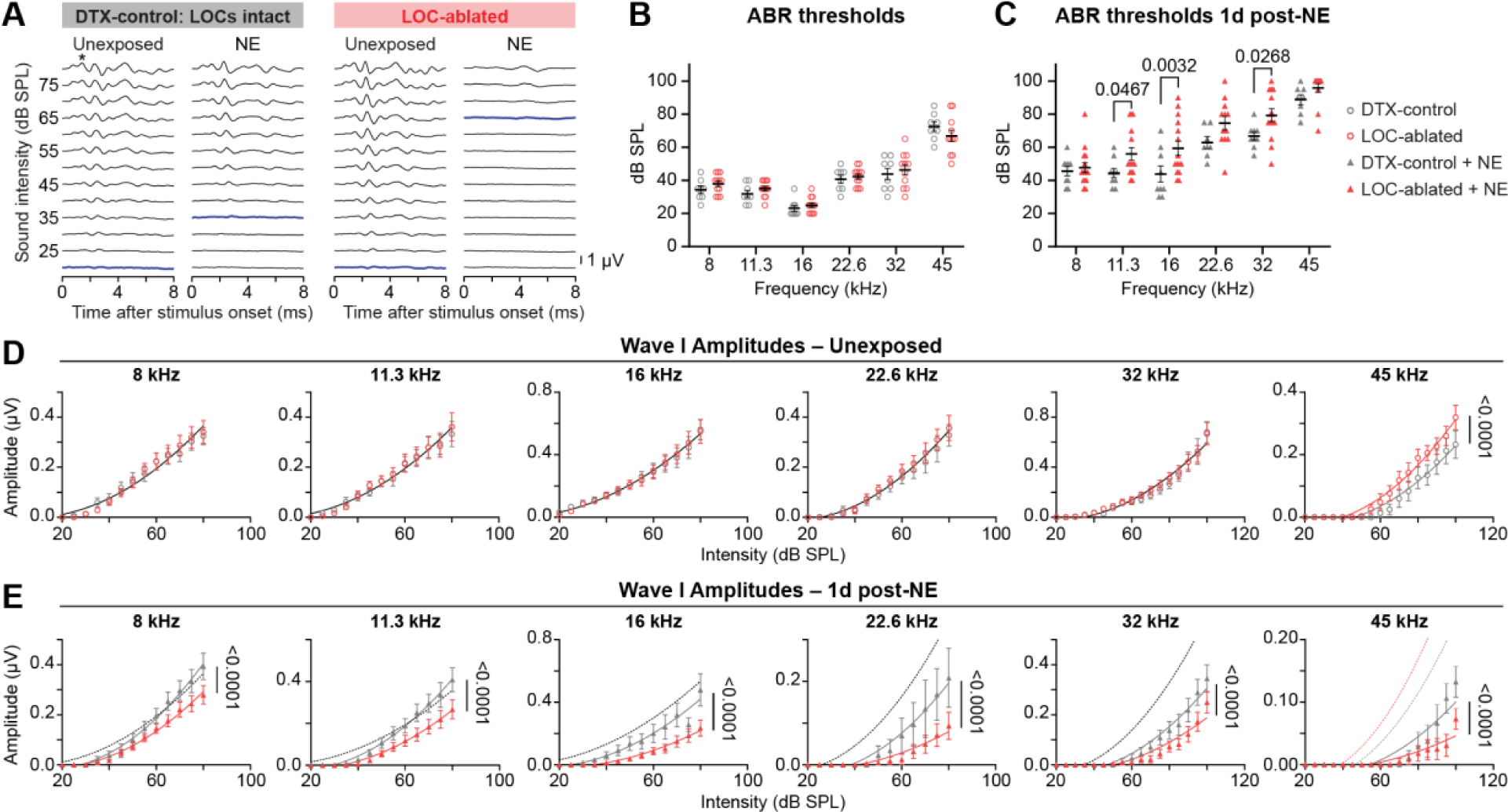
Animals lacking LOCs were more vulnerable to noise-induced functional deficits one day after noise exposure. **(A)** Example 16 kHz ABR waveforms from DTX-control mice or LOC-ablated mice. Example traces correspond to unexposed mice (left), or noise-exposed (NE) mice 1 day (1d) after NE (right). Blue lines denote ABR threshold. Asterisk marks the location of wave I at 80 dB SPL. (**B–C**) ABR thresholds in unexposed **(B)** and NE **(C)** mice. Two-way ANOVA with a post-hoc Tukey’s multiple comparisons test. Only significant two-way comparisons are labeled (*p* < 0.05). (**D-E**) Wave I amplitudes in unexposed **(D)** and NE mice **(E)** 1d after NE. Data were fit with a second order polynomial (quadratic) using a least-squares regression. Fits were compared using the extra sum-of-squares F-test. A solid black line indicates that one curve best fit both data sets (*p* > 0.05), whereas solid red (LOC-ablated) or solid gray (DTX-control) lines mean that each data set were best fit by different curves (*p* < 0.05). Dotted lines in **(E)** represent fits from panel **D**. Error bars, mean ± SEM. LOC ablated: unexposed, *N* = 11–12; NE, *N* = 13–15. DTX controls: unexposed, *N* = 8; NE, *N* = 7–9 mice.

### Reduced LOC input to the cochlea led to worse auditory recovery following acoustic trauma

Next, as LOC neurons are known to alter their transmitter expression in response to loud sound (**Fig. 1**) (5, 15, 33), we used acoustic trauma as a tool to reveal how this adaptive feature influences cochlear responses. To do so, we used a NE paradigm that induced temporary, recoverable shifts in the threshold at which sound is detectable (**Figs. 3B & S2**). This NE approach differs from the one in our sequencing experiments (**Fig. 1**), in which we used noise exposure levels that typically induce permanent deficits in auditory function (22). This more moderate NE paradigm allowed us to assess how loss of LOCs affected audition both shortly following the acoustic insult (1d post NE; **Fig 4**) and after a 2w recovery period post NE (**Fig. 5**). Mice with intact (DTX-control and SAL-control mice) or ablated LOCs were exposed to an 8–16 kHz broadband noise stimulus at 97.5 dB for 2h, at least 7d following the second injection of either DTX or SAL (**Fig. 3B**). Unexposed littermate controls in all three conditions were left in their home cages in the vivarium. We conducted ABRs on NE mice and unexposed littermates 1d after NE (**Fig. 4**) and again after a 2w recovery period (**Fig. 5**), when the threshold for sound detection returned to baseline in control animals (**Fig. S2C–D**).

**Figure 5.**
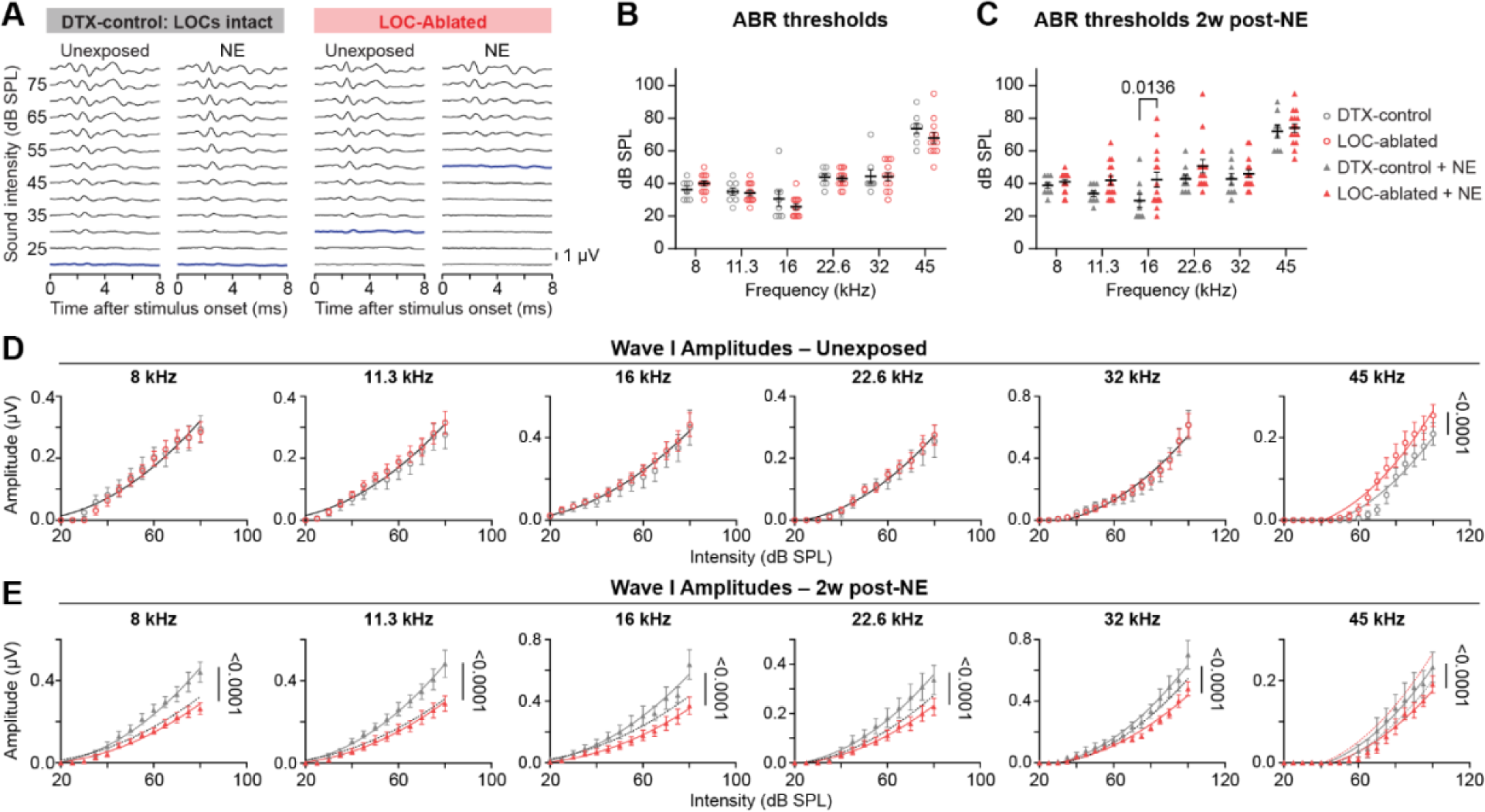
Animals lacking LOCs failed to fully recover from acoustic insult two weeks later. **(A)** Example 16 kHz ABR waveforms from DTX-control or LOC-ablated mice. Example traces correspond to unexposed mice (left), or 2 weeks (2w) after noise exposure (NE, right). Blue lines denote ABR threshold. **(B–C)** ABR thresholds in unexposed mice 2w after the first ABR **(B)** and NE mice 2w post-NE **(C)**. Two-way ANOVA with Tukey’s *post hoc* multiple comparisons test. Only significant two-way comparisons are labeled (*p* <0.05). (**D-E**) Wave I amplitudes in unexposed **(D)** and NE **(E)** mice. Data were fit with a second order polynomial (quadratic) using a least-squares regression. Fits were compared using the extra sum-of-squares F-test. A solid black line means that one curve best fit both data sets (*p* > 0.05), whereas solid red (LOC-ablated) or solid gray (DTX-control) lines mean that each data set was best fit by different curves (*p* < 0.05). Dotted lines in **E** represent fits from panel **D**. Error bars, mean ± SEM. LOC ablated: unexposed, *N* = 15 mice; NE, *N* = 12. DTX control: unexposed, *N* = 8–9; NE, *N* = 8.

As expected, 1d after NE, both LOC-ablated and DTX-control mice had elevated thresholds at frequencies ≥11.3 kHz, indicative of worsened auditory function when compared to unexposed animals (**Fig. S2A & C**). However, ABR thresholds were significantly higher in LOC-ablated animals than in DTX controls 1d after NE at 11.3, 16, and 32 kHz (**Fig. 4C**) and wave I amplitudes were lower at every frequency tested (**Fig. 4E**). Together, these results suggest that LOCs may help preserve cochlear sensitivity within a day after traumatic NE, either by preventing damage or by inducing a compensatory response in SGNs.

We next assessed auditory function in the same animals 2w after NE to determine how loss of LOC innervation affected recovery. At this time point, there continued to be no differences in ABR thresholds between unexposed DTX-control and unexposed LOC-ablated mice (**Fig. 5B**), nor any differences in wave I amplitudes, except at 45 kHz (**Fig. 5D**), similar to measurements from the ABR results two weeks prior (**Fig. 4D**). However, although ABR thresholds were fully recovered in NE DTX-control mice (**Fig. S2D**), elevated thresholds persisted at 16 kHz in NE LOC-ablated mice compared to NE-DTX controls (**Fig. 5A–C**) or to unexposed LOC-ablated animals (**Fig. S2B**). Additionally, 2w after NE, wave I amplitudes were significantly lower in LOC-ablated mice compared to DTX-controls at every frequency (**Fig. 5E**). Paradoxically, this result did not reflect worsened wave I amplitudes in LOC-ablated mice; rather, there was an overshoot in the amplitudes of NE DTX-control mice when compared to unexposed mice of either condition (**Fig. 5E**, dotted vs solid lines). This suggests that after NE in DTX-control animals, LOC neurons sensitized SGN excitability, perhaps as a means of restoring or maintaining auditory acuity to pre-NE levels, and this sensitization persisted for at least 2 weeks.

### Administration of diphtheria toxin, even in the absence of its receptor, led to loss of inner hair cells

Comparison between the two control conditions with intact LOCs, i.e., DTX-control and SAL-control mice, revealed that NE SAL-controls lacked the compensatory LOC-dependent response (**Fig. S3F, H**) we found in NE DTX-controls (**Figs. 4E, 5E**). This finding suggested that DTX might directly influence how the cochlea responds to acoustic trauma. Indeed, although DTX endocytosis should in theory be restricted to cells expressing DTR, our histological and functional data demonstrated that DTX alone was sufficient to damage IHCs (**Fig. S4**) and hinder peripheral auditory system function (**Fig. S3**), even in the absence of DTR. Micrographs of cochlear turns revealed stochastic loss of IHCs along the length of the cochlea (**Fig. S4A**). Regardless of NE status, the number of IHCs along a ∼100 μm length of the sensory epithelium at 8, 16, and 45 kHz was lower in mice that received DTX (**Fig. S4B, Table 2**). The differences in IHC number between mice that received DTX and those that received SAL were statistically significant (*p* < 0.05, Ordinary one-way ANOVA with Tukey’s *post hoc* multiple comparisons test), except for DTX-control mice at 8 kHz (**Fig. S4B**). There were no statistically significant differences in number of IHCs between either group of mice that received DTX, i.e., LOC-ablated and DTX-control mice (**Fig. S4B**), indicating that the ototoxic effect of DTX on IHCs did not require expression of DTR.

**Table 2.**
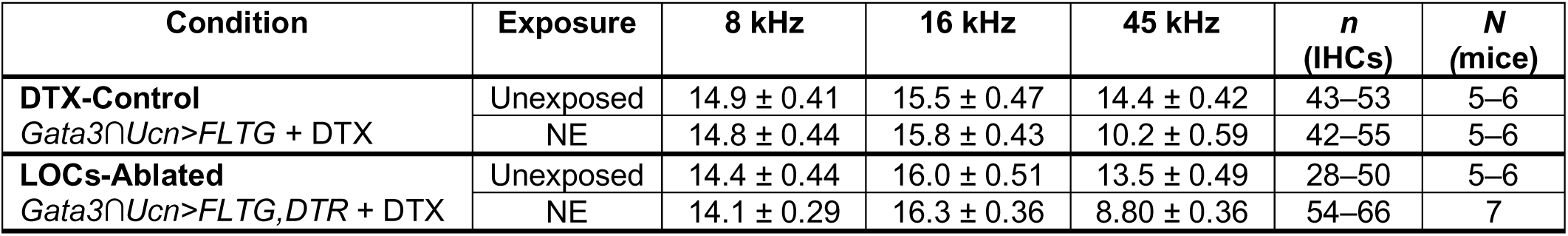
Number of Ctbp2+ puncta/IHC.

| Condition | Exposure | 8 kHz | 16 kHz | 45 kHz | *n* (IHCs) | *N* (mice) |
| --- | --- | --- | --- | --- | --- | --- |
| **DTX-Control**<br>*Gata3*∩*Ucn*>*FLTG* + DTX | Unexposed | 14.9 ± 0.41 | 15.5 ± 0.47 | 14.4 ± 0.42 | 43–53 | 5–6 |
|  | NE | 14.8 ± 0.44 | 15.8 ± 0.43 | 10.2 ± 0.59 | 42–55 | 5–6 |
| **LOCs-Ablated**<br>*Gata3*∩*Ucn*>*FLTG*, *DTR* + DTX | Unexposed | 14.4 ± 0.44 | 16.0 ± 0.51 | 13.5 ± 0.49 | 28–50 | 5–6 |
|  | NE | 14.1 ± 0.29 | 16.3 ± 0.36 | 8.80 ± 0.36 | 54–66 | 7 |

Prior reports showed that DTX ototoxicity led to elevated ABR thresholds when administered to mice <28d old, whereas mice that received DTX after that age had unaffected thresholds (40). Similarly, we found that ABR thresholds were similar between DTX-control and SAL-control mice at both the 1d and 2w time points in both unexposed and NE animals (**Fig. S3A–D**). In contrast, when we compared ABR wave I amplitudes, we found that DTX-control mice had lower amplitudes than SAL controls (**Fig. S3E–G**). This difference in amplitudes between DTX and SAL controls was found at both time points, regardless of whether they received NE, and was statistically significant (*p* < 0.05, sum of squares F-test) at most frequencies tested. These data indicate that the DTX-driven loss of IHCs reduced SGN responses to sound.

We also considered the possibility that either DTX itself or the loss of LOCs might alter OHC function and hence indirectly lead to dampened ABR responses. Similarly to a prior study (41), we observed no gross differences in the number of OHCs in DTX-treated animals. To further assess cochlear function, we recorded distortion product otoacoustic emissions (DPOAEs), which are sounds made by the inner ear that reflect the function of OHCs, which enhance the detection of sound in the cochlea (10, 12, 42). Unexposed DTX-control, SAL-control, and LOC-ablated mice had similar DPOAE thresholds at all frequencies (**Fig. S5A, C**). As expected, 1d after NE, all three groups had elevated DPOAE thresholds at frequencies ≥22.6 kHz (**Fig. S6A, C, E**), which recovered completely 2w later (**Fig. S6B, D, F**), except for 22.6 kHz in LOC-ablated mice, which remained elevated relative to unexposed LOC-ablated mice (**Fig. S6D**). However, when comparing among all NE mice, DPOAE thresholds did not significantly differ between DTX-control, SAL-control, or LOC-ablated mice either 1d (**Fig. S5B**) or 2w after NE (**Fig. S5D**), consistent with the prediction that ablation of LOC neurons had only a minor effect on OHC function. Instead, the presence of significantly diminished ABR responses in NE LOC-ablated animals when compared to NE DTX-controls are best explained by changes in SGN activity.

Taken together, these results highlight the importance of analyzing multiple features of cochlear function—including DPOAE thresholds, ABR thresholds, and ABR wave I amplitudes— to gauge auditory sensitivity, as well as of comparing SAL-controls to animals that received DTX but lack DTR when conducting DTX ablation experiments.

### Loss of LOCs did not alter the number of presynaptic puncta at the SGN:IHC synapse

One way that LOCs may preserve cochlear function following noise exposure is by minimizing SGN synaptopathy, which is a common mechanism underlying permanent loss of function following a traumatic noise exposure (18, 43–45). To test this idea, we examined putative synaptic connections between Type I SGNs and IHCs 2w after NE in LOC-ablated and DTX-control mice by analyzing the number of presynaptic ribbons on IHCs (**Fig. 6**). As nearly all IHC pre-synaptic ribbons continue to be apposed to postsynaptic glutamate receptors within a week after a temporary threshold shift (25, 46), analysis of Ctbp2+ presynaptic puncta is a reliable readout of synapse number.

**Figure 6.**
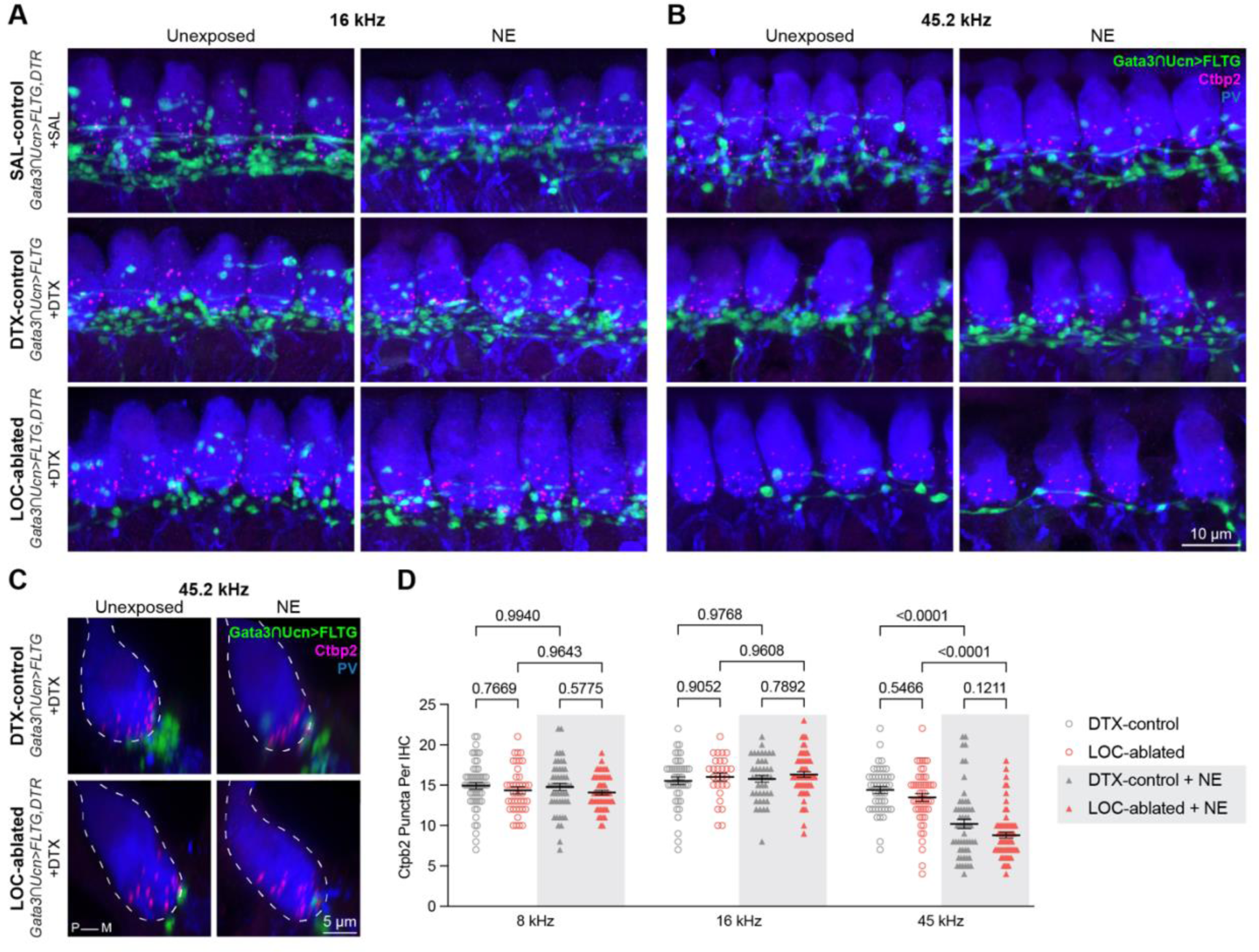
Number of presynaptic ribbons was unaffected by LOC ablation. **(A–B)** Representative images of presynaptic ribbons (Ctbp2, magenta), LOCs (Gata3∩ Ucn>FLTG, green), and IHCs and SGN peripheral processes (PV, blue) in unexposed and noise exposed (NE) mice in all three experimental groups (SAL-control, DTX-control, and LOC-ablated) at cochlear frequency regions 16 kHz **(A)** and 45.2 kHz **(B)**, 15–20 days after NE (i.e., 1–5 days after second ABR). **(C)** Orthogonal 3D snapshots of individual IHCs from unexposed and NE DTX-control and LOC-ablated cochlea at 45.2 kHz. IHCs are outlined in dotted lines. **(D)** Number of Ctbp2 puncta per IHC at three frequencies. There were no statistically significant differences in the number of Ctbp2 ribbons between LOC-ablated and DTX-control cochlea at any frequency analyzed. At 45 kHz, there were fewer ribbon puncta in NE cochlea, but no statistically significant differences between LOC-ablated and DTX-control mice. Error bars, mean ± SEM. Ordinary one-way ANOVA with Tukey’s *post hoc* multiple comparisons test. See **Table 2** for means, SEM, and sample sizes. DTX = diphtheria toxin; SAL = saline; NE = noise exposure; P = pillar; M = modiolar; PV = Parvalbumin

Exposure to an 8–16 kHz band of noise typically induces synapse loss in higher (32 and 45 kHz) frequency regions of the cochlea (18). As expected, NE caused a loss of pre-synaptic puncta in the 45 kHz region of the cochlea in both DTX-control and LOC-ablated mice when compared to unexposed animals (**Fig. 6B–D**). The degree of the effect was similar regardless of the presence of LOCs, with a ∼29% reduction in ribbons per IHC in DTX-control mice (presynaptic puncta/IHC: unexposed 14 ± 0.42 vs NE 10 ± 0.59, *p* < 0.0001, means ± SEM, one-way ANOVA) and a ∼32% reduction in LOC-ablated mice (unexposed 13 ± 0.49 vs NE 8.8 ± 0.36, *p* < 0.0001, means ± SEM, one-way ANOVA). There were no significant differences in pre-synaptic puncta number between LOC-ablated and DTX-control mice in either the unexposed or NE conditions at this frequency (**Fig. 6B–D**, **Table 3**). Thus, loss of LOCs did not exacerbate synaptic pathology in high-frequency regions of the cochlea.

**Table 3.** Number of IHCs/100 μm.

| Condition | Exposure | 8 kHz | 16 kHz | 45 kHz | *N* (mice) |
| --- | --- | --- | --- | --- | --- |
| **SAL-Control**<br>*Gata3*∩*Ucn*>*FLTG*, *DTR* + SAL | Unexposed | 11 ± 0.14 | 12 ± 0.36 | 12 ± 0.22 | 6–7 |
|  | NE | 11 ± 0.37 | 13 ± 0.22 | 12 ± 0.48 | 6 |
| **DTX-Control**<br>*Gata3*∩*Ucn*>*FLTG* + DTX | Unexposed | 8.8 ± 0.48 | 8.6 ± 1.3 | 6.8 ± 0.66 | 5–6 |
|  | NE | 8.5 ± 0.56 | 8.4 ± 0.51 | 9.4 ± 0.75 | 5–6 |
| **LOCs-Ablated**<br>*Gata3*∩*Ucn*>*FLTG*, *DTR* + DTX | Unexposed | 6.8 ± 0.65 | 6.6 ± 1.1 | 8.3 ± 0.67 | 5–6 |
|  | NE | 7.7 ± 0.81 | 8.0 ± 0.82 | 9.4 ± 0.43 | 7 |

Since ABR thresholds to 16 kHz tones were significantly elevated in LOC-ablated animals 2w post-NE (**Fig. 5C**) and ABR wave I amplitudes were significantly smaller in LOC-ablated animals at both 8 and 16 kHz (**Fig. 5E**), we also quantified pre-synaptic puncta in these regions of the cochlea. Following the typical pattern of pathology, there was no loss of pre-synaptic puncta from the 8 or 16 kHz region of DTX-control mice after noise exposure (unexposed: 8 kHz: 15 ± 0.41, 16 kHz: 16 ± 0.47; NE: 8 kHz: 15 ± 0.44, 16 kHz: 16 ± 0.43; means ± SEM, *n* = 28–56 IHCs, *N* = 5–7 mice of both sexes; **Fig. 6A, D**). Notably, this was also true in LOC-ablated animals, which had the same number of pre-synaptic puncta in both the unexposed and NE conditions (unexposed: 8 kHz: 14 ± 0.44, 16 kHz: 16 ± 0.51; NE: 8 kHz: 14 ± 0.29, 16 kHz: 16 ± 0.36). Thus, any difference in ABR responses between mice with or without intact LOCs at 8, 16, or 45 kHZ frequencies is not due to enhanced loss of pre-synaptic ribbons. Altogether, these results show that a reduction of LOC innervation in the cochlea had no apparent effect on the number of presynaptic puncta either at baseline or 2w following traumatic NE. These findings support the idea that LOCs preserve cochlear function by directly modulating SGN activity rather than by preventing noise-induced synapse loss.

## DISCUSSION

LOCs have long been implicated in providing centrally mediated feedback to the cochlea that can alter how animals detect and encode sound stimuli. Here, we show that LOC-mediated feedback modulates how the cochlea functions after exposure to traumatic levels of noise. We developed genetic tools that allow selective access to LOCs and assayed cochlear function and anatomy before and after noise exposure (NE). Our evidence suggests that LOCs can enhance the cochlear response to sound after NE, likely by increasing SGN excitability for days to weeks after exposure. In support of this idea, LOCs transiently upregulate a suite of transcripts by 1d after NE, including those of the neuropeptides *Calca*, *Ucn*, and *Npy*. Additionally, we found no evidence that poor auditory responses were due to decreased numbers of putative SGN:IHC synapses in mice without an intact LOC system. Rather, we propose that, in response to loud sounds, LOCs play a compensatory role and preserve auditory function by offsetting the excitability of SGNs to counteract loss of function resulting from damage to the cochlear sensory epithelium.

Housed in the brainstem, both major groups of the olivocochlear efferents are poised to directly modulate cochlear function based both on the immediate needs of the animal and the external auditory environment. A large body of evidence indicates that MOCs inhibit OHC electromotility, thereby decreasing overactivation of IHCs (10, 23–25, 27, 47–50). Such a feedback mechanism can offer immediate protection from noise-induced hearing loss, although there is evidence that alternative mechanisms could underlie MOC-mediated protection (51, 52). The contributions of LOCs, on the other hand, have been less clear (14, 25, 27, 53, 54). Our data show that LOCs also influence cochlear function, but through a different mechanism and on a different time scale than MOCs. Specifically, the simplest explanation for our molecular, physiological, and anatomical data is that LOCs directly modulate SGN activity, both in normal hearing environments and in the days to weeks following noise exposure. In support of this idea, ABR wave I amplitudes, which reflect the number and synchrony of SGNs activated by sound, were decreased in LOC-ablated animals compared to DTX-control mice two weeks after NE (**Fig. 5E**). In contrast to what occurs when the MOC system is disabled, OHC function was essentially preserved in LOC-ablated animals, as determined by measuring DPOAEs (**Fig. S5–S6**). Additionally, the number of presynaptic puncta in IHCs at the SGN:IHC synapse did not differ between LOC-ablated and DTX-control mice, either in baseline conditions or following NE (**Fig. 6**). Since cochlear synaptopathy is reliably accompanied by pre-synaptic ribbon loss (18), these data argue against the hypothesis that LOCs protect SGN synapses from NE and point instead toward direct modulation of SGN activity. Indeed, it is well established that LOC axons terminate on SGN peripheral processes (7, 55). Additionally, we find that LOCs—and not MOCs—show increased expression of multiple neuropeptide signaling molecules after NE that lasts for at least a week, indicating one possible mechanism by which LOC signaling onto SGNs may be altered by NE.

Consistent with their heterogeneous and dynamic properties, LOCs appear to have a variety of effects on cochlear function depending on the context. For instance, in baseline conditions, LOCs appeared to inhibit SGN activation in high-frequency regions of the cochlea, as evidenced by increased wave I amplitudes in LOC-ablated animals compared to DTX controls (**Figs. 4D & 5D**). On the other hand, after NE, the opposite effect was observed: LOC-ablated animals showed worse auditory responsiveness than DTX controls in both the short (1d) and long (2w) term following NE (**Figs. 4E & 5E**). However, the difference in auditory function between DTX-control and LOC-ablated animals two weeks post-NE did not stem from lasting depression of wave I amplitudes in the LOC-ablated mice. Instead, NE DTX-control mice had elevated amplitudes relative not only to NE LOC-ablated mice but also to unexposed DTX-control and unexposed LOC-ablated mice across many frequencies (i.e., 8–22.6 kHz), as seen by comparing the amplitudes of NE mice (solid lines in **Fig. 5E**) to those of unexposed mice (dotted lines in **Fig. 5E**, based on the line of best fit in **Fig. 5D**). This is an unexpected result, as enhanced wave I amplitudes have not been previously reported in noise-exposed mice, consistent with our findings in SAL-control mice. One possible explanation is that this LOC-dependent effect is only present in the context of a damaged cochlea. In fact, all animals treated with DTX showed significant loss of IHCs and hence decreased cochlear function, independent of noise exposure. Something about this condition appears to uncover a role for LOCs upon noise exposure. Because the only difference between LOC-ablated mice and DTX-controls is the presence of LOCs, our data suggest that loss of IHCs enhances LOC-dependent compensation of SGN activity after NE. This result is consistent with single-fiber recordings demonstrating hyperexcitability of SGNs two weeks after a synaptopathic NE (56). Although wave I amplitudes were not enhanced in this study, it is important to note that the IHCs were not damaged at the time of noise exposure as they were in the DTX-controls in the present study. It has been suggested that LOCs modulate SGN sensitivity in response to changing environmental contexts and perhaps to internal states (3, 28, 57). Our data align with this interpretation: rather than directly protecting against cochlear damage, LOCs appear to increase SGN excitability over an extended period of days to weeks, perhaps as a means of enhancing sound detection in times of stress or cochlear damage.

The identity of the upstream circuits that lead to LOC activation and consequent changes in gene expression following NE are currently unknown. Although we show that an intense sound stimulus is sufficient to induce transcriptional changes in LOCs (**Fig. 1**), it is unclear whether the sound stimulus itself, the stress of noise exposure, or some combination of the two is driving the inputs that cause these gene-expression changes. Indeed, many of the neuropeptides that are increased in LOCs following NE (**Fig. 1I**) are associated with stress-response pathways (58–61). Moreover, previous work has found that stressful events—such as sham surgery or physical restraint—are sufficient to improve auditory responses after NE, perhaps by recruiting similar changes in LOC signaling to those seen here (28, 62). With the availability of genetic access to LOCs afforded by *Gata3^FlpO^;Ucn^Cre^* mice, future studies can further examine the flexibility of LOC functions in diverse contexts.

Another piece of the puzzle is the fact that high-frequency regions, including 45 kHz, are densely innervated by a transcriptionally distinct subset of LOCs, the LOC2s, which consistently express high levels of neuropeptides (5). LOC2s are identifiable at birth (before hearing onset) based on restricted expression of neuropeptide Y (NPY) in the medial wing of the LSO. This group of LOCs innervate high-frequency regions of the cochlea more densely than lower frequencies (5), including 45 kHz, where LOCs dampened SGN excitability at baseline (**Figs. 4D, 5D**); however, it is unclear whether these two phenomena are functionally related. We and others have shown previously that LOCs can upregulate a suite of neuropeptide and neurotransmitter proteins after NE in both guinea pigs and mice (5, 15, 33). Here, we show that NE led to transient transcriptional changes in LOCs 1d after NE, which waned or altogether disappeared 7d later (**Fig. 1**). Despite the transient nature of these transcriptional changes, altered expression of TH and neuropeptide proteins persists for at least a week after NE (5, 15), likely reflecting the fact that protein half-lives are much longer than that of their mRNA templates (63, 64).

A greater fraction of LOCs clustered with LOC2s 1d after NE relative to unexposed littermates or NE mice 7d after the exposure (**Fig. 1K**), driven by elevated expression of more LOC2-like features, including transcripts for the neuropeptides *Calca*, *Npy*, and *Ucn* (**Fig. 1I**). However, this binary classification scheme belies a more nuanced reality: as with cellular subtypes identified elsewhere in the brain (65–68), LOC subtypes appear to exist along a spatially defined continuum, with a gradient of gene expression blending LOC2s into LOC1s along the medial–lateral axis of the LSO (5). We have previously shown that this gradient persists at the protein level after NE, even as the fraction of peptide-expressing LOCs increases (5). Determining the gene-regulatory programs that establish this expression gradient and examining how these programs are altered by LOC activity constitutes a rich avenue for future study. Although we were unable to detect changes in *Th* expression in our sequencing data—likely due to the low expression of *Th* transcripts—it has previously been shown that induced TH expression in LOCs does not follow the same medial–lateral gradient as do neuropeptides (5, 15). Thus, LOC neurons may possess multiple activity-dependent gene-expression programs that drive altered transcription of these various signaling molecules.

The effects of LOC signals in the cochlea may be similarly multifaceted. Although dopamine appears to inhibit SGN activity (15, 31), acetylcholine and CGRP seems to increase afferent excitability (69–75). Moreover, analysis of dopamine receptor mutant mouse lines indicates that even the same molecule can have opposing effects on vulnerability to NE depending on what receptor is involved (76). The contributions of other peptides expressed by LOCs are less clear, although transcripts for the NPY receptor *Npy1r* were detected in a subset of Type I SGNs (77–79); multiple cell types in the cochlea also appear to express the Ucn receptors *Crhr1* and *Crhr2*, including SGNs, supporting cells, and hair cells (77–82). Global knock-out of the Ucn receptor *Crhr2* leads to a decrease in ABR thresholds, suggesting that Ucn may inhibit auditory activity at baseline (80). However, germline deletion of *Ucn* or the Ucn receptor *Crhr1* causes an elevation in both ABR and DPOAE thresholds (81, 83), indicating that Ucn and CRH peptides may have multiple effects in the cochlea, including effects on cochlear cells other than SGNs. The observation that signaling molecules expressed by LOCs can have seemingly opposing effects on SGN activity fits with previous *in vivo* recordings following indirect LOC activation (14), as well as our own finding that LOCs appear to repress activity in high-frequency regions at baseline (**Figs. 4D & 5D**) while increasing excitability of SGNs after sound exposure in DTX-damaged cochleae (**Figs. 4E & 5E**). Whether this NE- and LOC-dependent increase in SGN excitability is a direct consequence of altered transmitter expression in LOCs remains to be determined.

Our experiments likely disrupt all of these disparate signaling events to the same degree. Consistent with the broad expression of Ucn during development, *Ucn^Cre^* drove expression of tdTomato in most LOCs, as identified by staining for ChAT (**Fig. 2E**). Further, in the LOC-ablated animals, we observed loss of LOCs throughout the LSO, both in the medial wing where LOC2s reside and more laterally where LOC1s predominate. As a result, there are large swaths of the cochlea that are completely devoid of LOC innervation (**Fig. 3E**), making it highly unlikely that a specific subset of LOCs is resistant to DTX-induced ablation. Finally, although it is likely that the remaining LOCs include some that are dopaminergic—either constitutively or after NE—our data indicate that this population is not sufficient to fully preserve auditory function. Thus, while our findings highlight an important role for the LOCs in controlling cochlear function overall, much more work is needed to decipher the relative contributions of each signaling system, especially given the fact that LOCs alter their transmitter profile in a dynamic fashion.

Within the cochlea, LOCs constitute one part of a larger tapestry of modulatory cells, including adrenergic sympathetic neurons and peptide-expressing glial and supporting cells. Several of the peptides upregulated by LOCs after NE, including CGRP and NPY, are also expressed by other cochlear cell types; similarly, some supporting cells appear to express CRH, which binds to the same receptors as Ucn (84, 85). Many of these supporting cells also express neuropeptide receptors, suggesting that they may be targets of peptidergic modulation as well as an additional source of neuropeptides (81, 82, 86). Immune cells within the cochlea may also be substrates of peptidergic signals, as the composition of cochlear immune cells is altered by sound exposure (86, 87) and neuropeptides are known to be immune modulators (88). Because neuropeptides can diffuse across large areas—with signaling distances of ∼100 µm measured *in vivo*—peptide-expressing cells in the ear may exert widespread effects on cochlear physiology even in the absence of direct synaptic contacts (89–91). The multifaceted interactions between these various cell types and their ultimate effects on auditory function remain an important open question.

## ACKNOWLEDGEMENTS

We are grateful to Lin Wu and the Harvard Genome Modification Facility for assistance with creating the *Ucn^Cre^*and *Gata3^FlpO^* mouse lines. We thank Qiufu Ma for providing the *Mapt^ds-DTR^* mouse line and Susan Dymecki for providing *Rosa26^FSF-tdTomato^* animals. Services and guidance from the Harvard Department of Immunology Flow Cytometry Core, Biopolymers Facility, and Bauer Core Facility were valuable for execution of sequencing experiments. Initial transcriptomic data processing was performed on the O2 High Performance Compute Cluster, supported by the Research Computing Group at Harvard Medical School. We thank the Neurobiology Imaging Facility for the use of the Olympus VS120 slide scanner. Kiera Grierson, Lorna Wu, and Sadie Schlabach assisted with ABRs; Cheuk Wong, Mackenzie Hunt, and Sarah Yoder provided histology assistance. We thank Katelyn Comeau Boulanger and Sydney Maeker for assistance and input on the project. We are grateful to M. Charles Liberman for advice and input throughout the project and for valuable feedback on the manuscript. This work was supported by R01 DC015974 (LVG), a Blavatnik Sensory Disorders Research Award (LVG), F32 DC019009 (AAS), and an HHMI Hanna H. Gray Fellowship (GER).

## AUTHOR CONTRIBUTIONS

**Conceptualization**: MMF, GER, AAS, LVG; **Formal analysis**: MMF, GER, AAS; **Investigation**: MMF, GER, AAS; **Writing**: MMF, GER, AAS, LVG; **Visualization**: MMF, MMF, AAS; **Supervision**: LVG; **Funding acquisition**: LVG, GER, AAS. MMF, GER, and AAS contributed equally; the order of authorship was determined by rolling a D20 die.

## DECLARATION OF INTERESTS

The authors declare no competing interests.

## RESOURCE AVAILABILITY

Sequencing data generated in this paper will be deposited on GEO and gEAR prior to publication.

## LEAD CONTACT

Further information and requests for resources and reagents should be directed to and will be fulfilled by the lead contact, Lisa Goodrich.

## SUPPLEMENTAL INFORMATION

**Figure S1.**
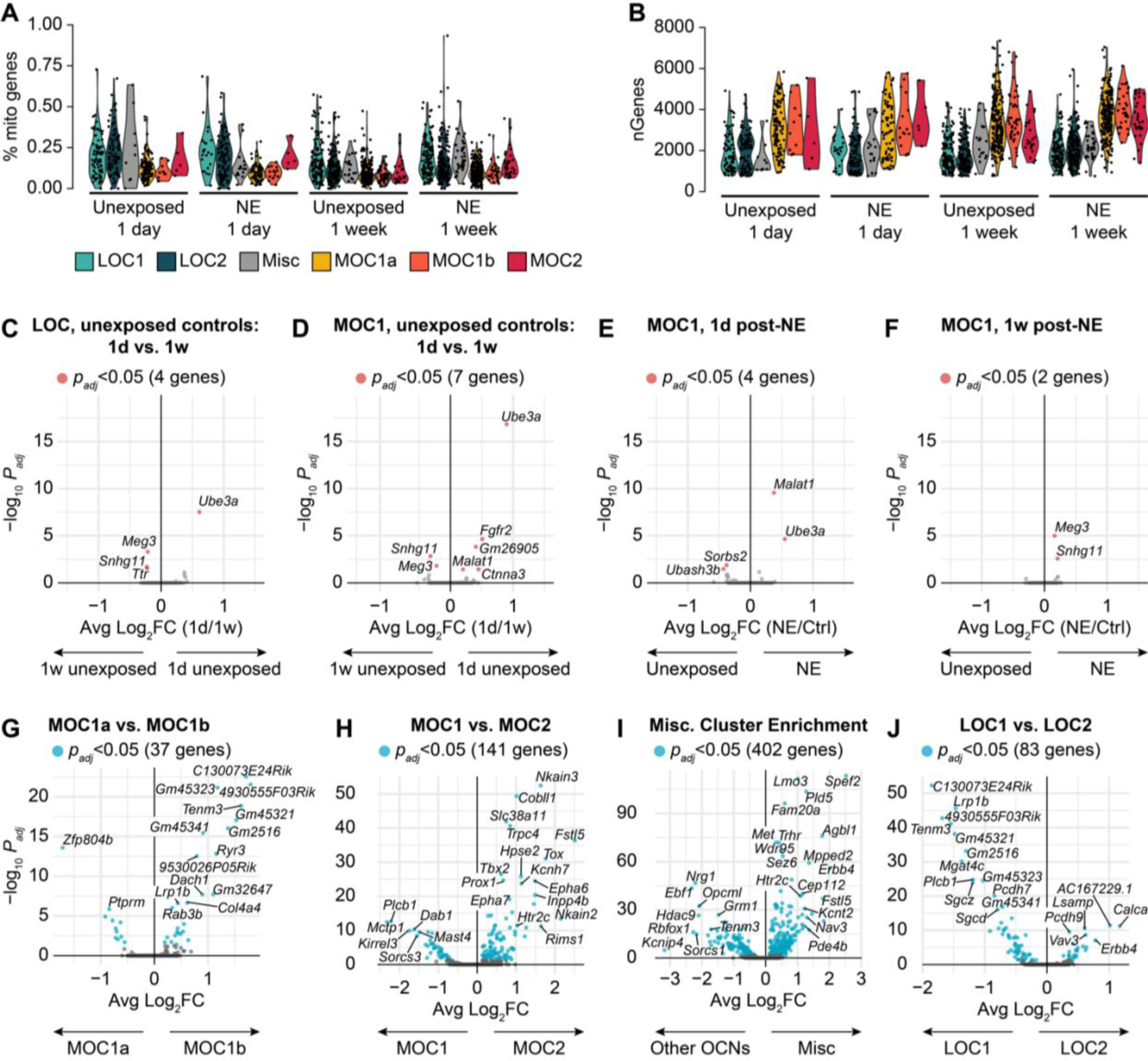
Additional single-nucleus sequencing quality-control and differential expression analysis. **(A, B)** Quality-control metrics of single-nucleus sequencing data. Cluster differences were not driven primarily by variability in technical noise such as cell health—as indicated by the fraction of mitochondrially encoded genes **(A)**—or the number of unique genes detected per cell **(B)**. **(C, D)** Variations in gene expression between 1 day (1d) and 1 week (1w) unexposed control LOC **(C)** and MOC **(D)** samples. In both MOCs and LOCs, a small number of genes were differentially expressed between unexposed littermate control samples collected at different time points, indicating the presence of some batch effects. **(E, F)** There were few differentially expressed genes in MOCs at 1d and 1w after exposure, which were largely explained by variability between control samples **(D)**. (**G–J**) Violin plots denoting differentially expressed genes between cell-type clusters from unexposed control animals. **(C–J)** Wilcoxon rank-sum test with Bonferroni *post-hoc* adjustment.

**Figure S2:**
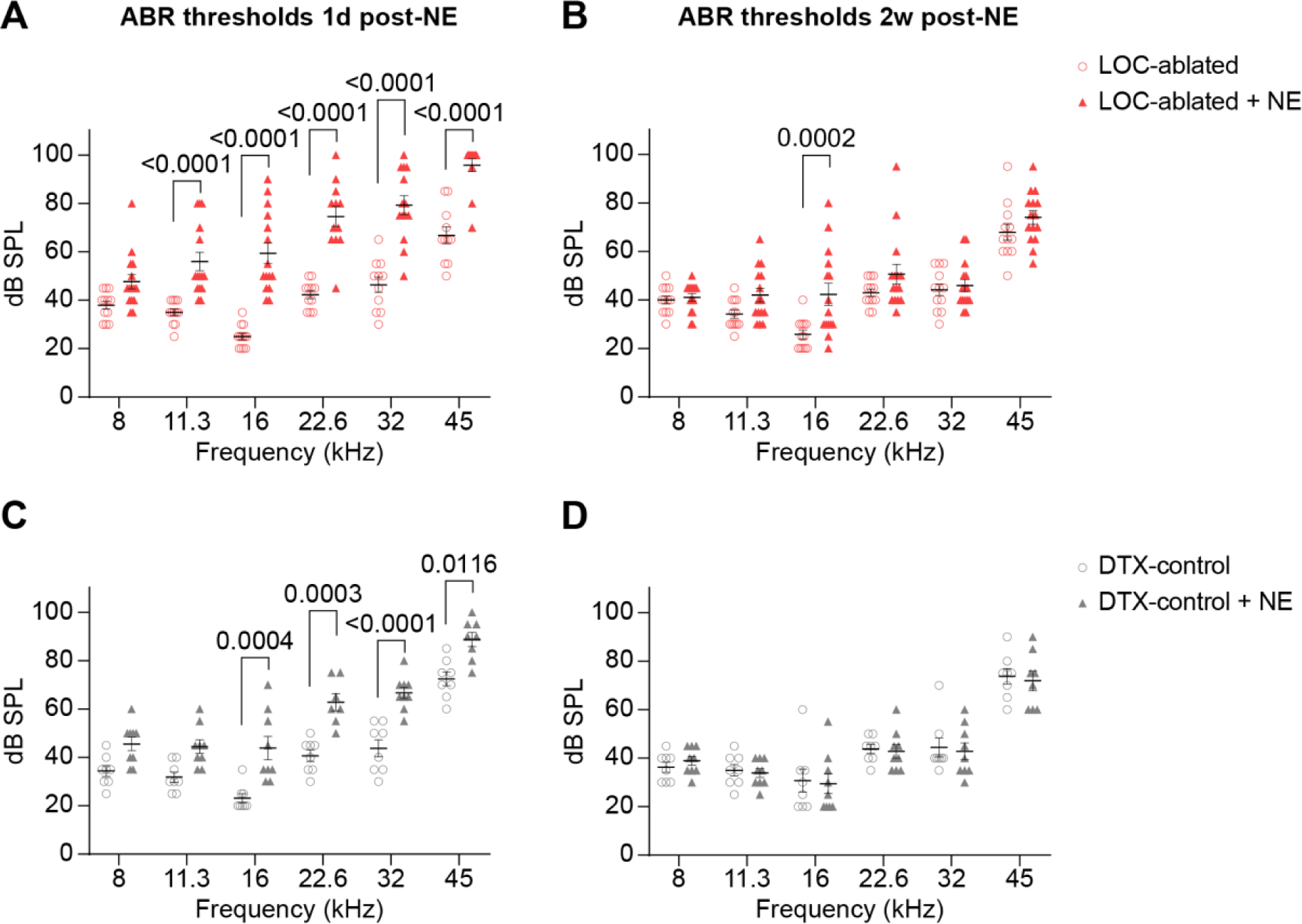
NE paradigm caused a temporary shift in ABR thresholds. **(A–B)** ABR thresholds in LOC-ablated mice that were unexposed (open red circles) or noise exposed (NE, red triangles) at 1 day (1d) or 2 weeks (2w) after NE. NE animals had elevated thresholds 1d after NE **(A)**, which recovered at 2w, except at 16 kHz **(B)**. **(C–D)** ABR thresholds in DTX-control mice that were unexposed (open gray circles) or NE (gray triangles) at 1d **(C)** or 2w **(D)** after NE. NE animals had elevated thresholds 1d after NE **(C)**, which recovered at 2w **(D)**. Error bars, mean ± SEM. Two-way ANOVA with a *post hoc* Tukey’s multiple comparisons test. Only significant two-way comparisons are labeled (*p* < 0.05). Sample sizes are the same as in Figs. 4 and 5.

**Figure S3.**
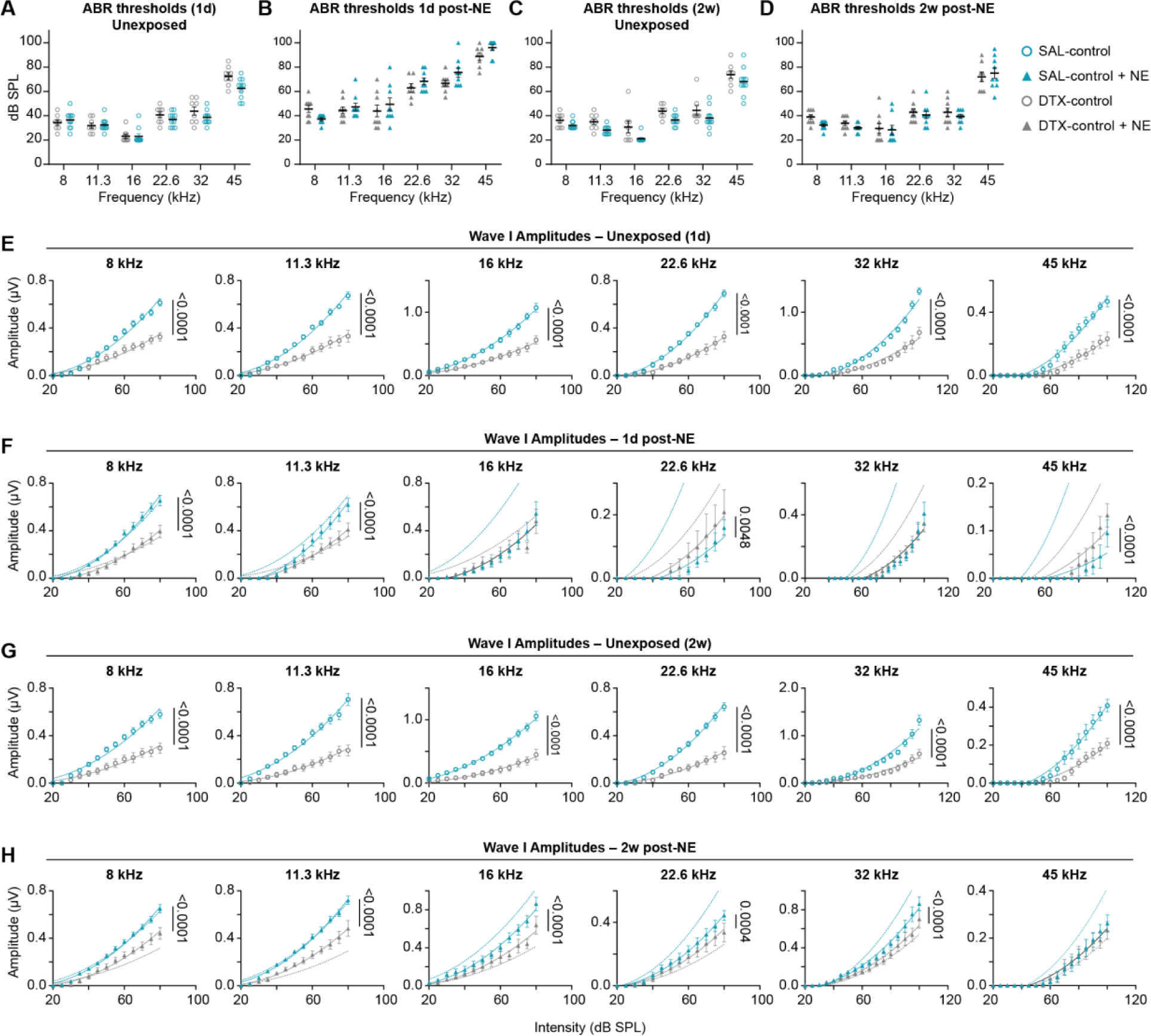
Diphtheria toxin alone worsened auditory function. **(A–D)** ABR thresholds in SAL-control (blue) and DTX-control (gray) mice. **(A–B)** Thresholds in unexposed **(A)** and noise exposed (NE) **(B)** mice 1 day (1d) after NE. **(C–D)** Thresholds in unexposed **(C)** and NE **(D)** mice 2 weeks (2w) after NE. Two-way ANOVA with a *post hoc* Tukey’s multiple comparisons test. Only significant two-way comparisons are labeled (*p* < 0.05). **(E–H)** Wave I amplitudes in SAL-control (blue) and DTX-control (gray) mice. **(E–F)** Amplitudes from unexposed **(E)** and NE **(D)** mice 1d after NE. **(G–H)** Amplitudes in unexposed **(G)** and NE **(H)** mice 2w after NE. Amplitude data were fit with a second order polynomial (quadratic) using a least-squares regression **(E, G)**. Fits were compared using the extra sum-of-squares F-test. A solid black line means that one curve best fit both data sets (*p* > 0.05), whereas solid blue (SAL-control) or solid gray (DTX-control) lines mean that each data set was best fit by different curves (*p* < 0.05). Dotted lines in **(F)** and **(H)** represent fits from **(E)** and **(G)**, respectively. Error bars, mean ± SEM. Sample sizes for DTX-control mice are the same as Figs. 4 and 5. SAL-control: unexposed, *N* = 10; NE, *N* = 9.

**Figure S4.**
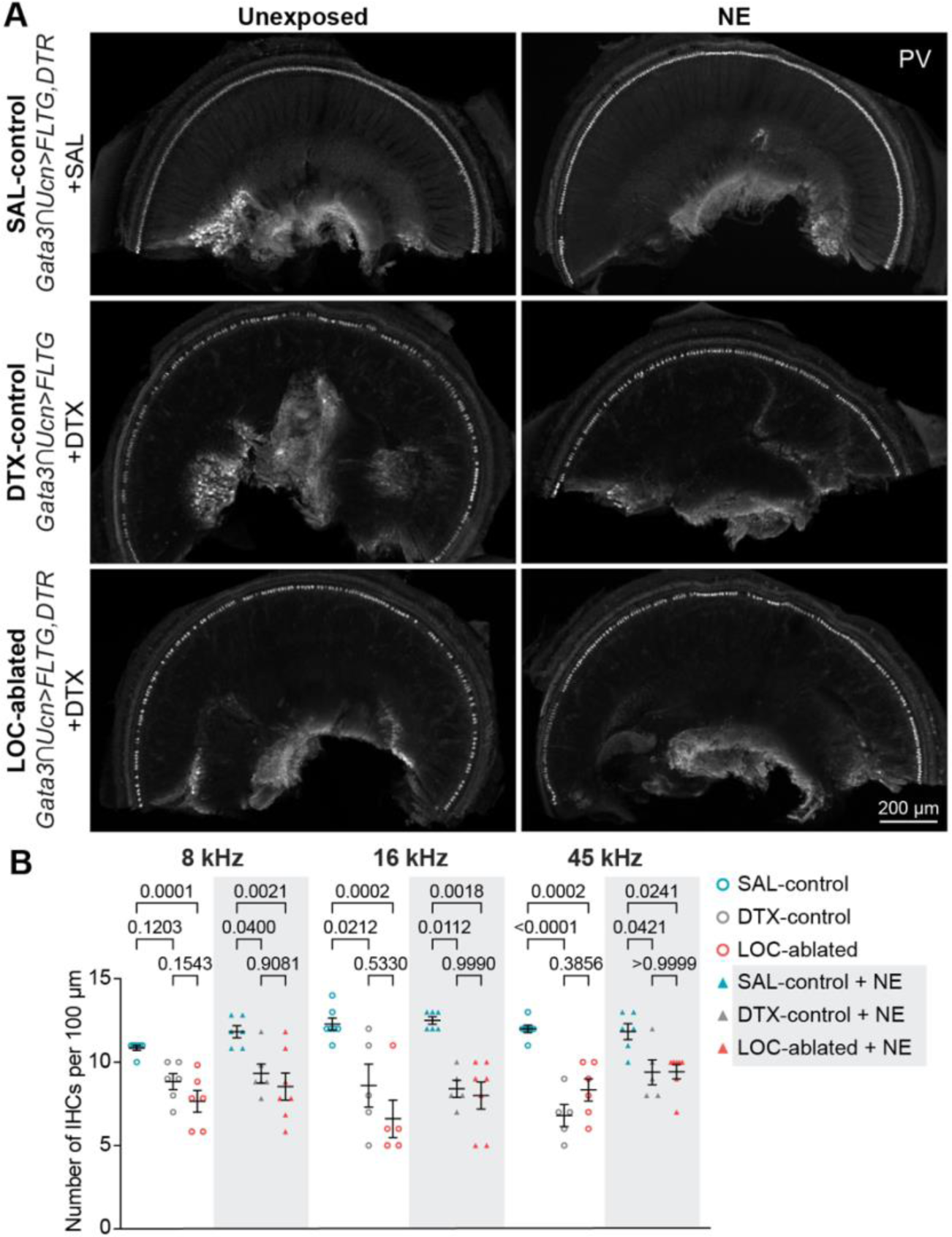
Diphtheria toxin is ototoxic to inner hair cells. **(A)** Micrographs of wholemounted cochlear base turns with IHCs labeled by parvalbumin (PV). Stochastic loss of IHCs is evident in mice that received DTX (i.e., DTX-control and LOC-ablated) compared with mice that did not receive DTX (SAL-control, top row). **(B)** DTX significantly reduced the number of IHCs along a 100 μm length of cochlea at three frequency regions, compared to mice that received SAL. There were no differences in the number of IHCs between mice that received DTX (i.e., DTX-control and LOC-ablated). Error bars, mean ± SEM. Ordinary one-way ANOVA with Tukey’s *post hoc* multiple comparisons test. *N* = 5–7 mice of both sexes in each condition.

**Figure S5:**
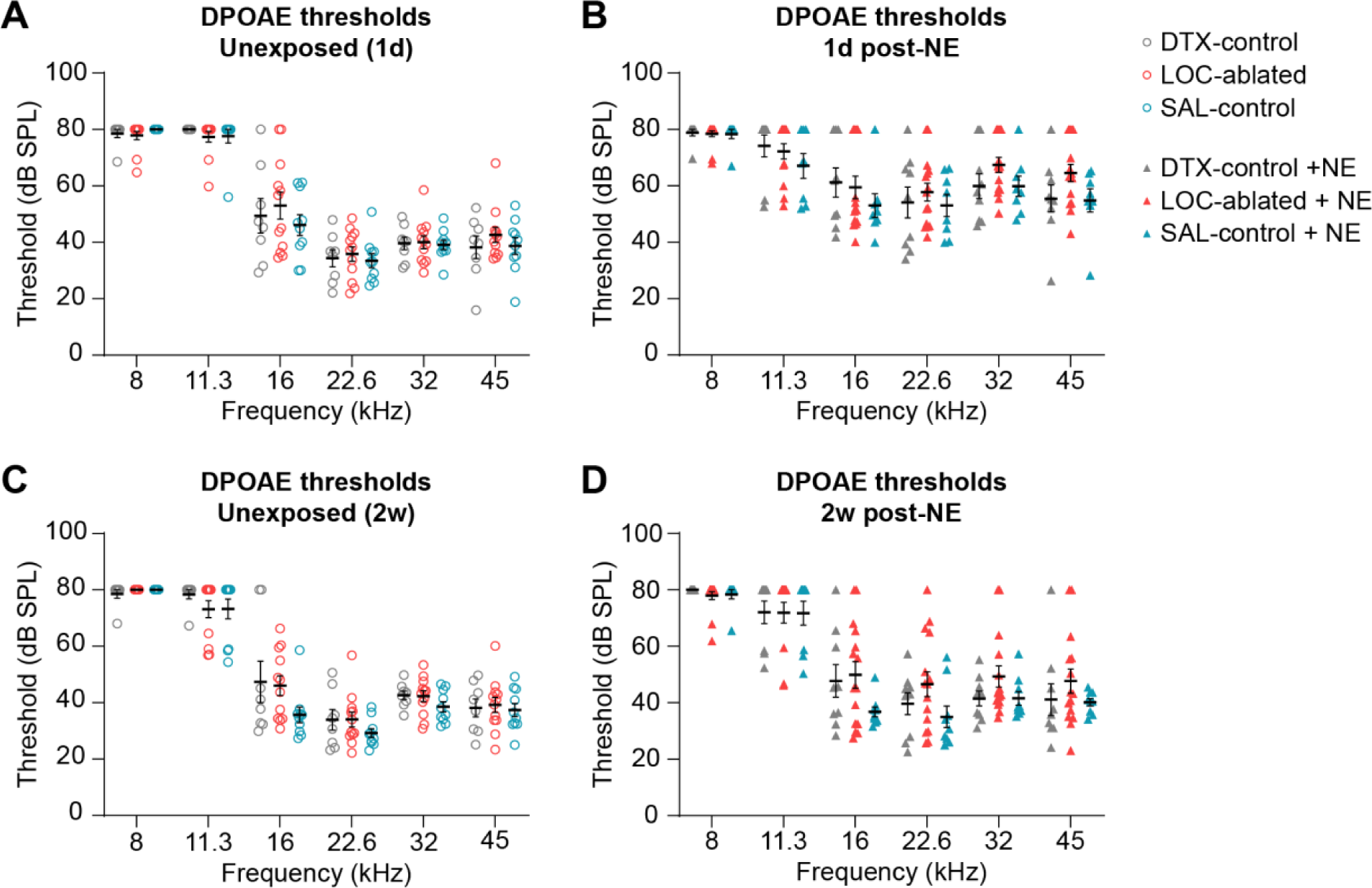
Outer hair cell function did not differ between mice with or without LOCs. **(A–B)** DPOAE thresholds in unexposed **(A)** and noise exposed (NE) mice **(B)** 1 day (1d) after NE. **(C–D)** DPOAE thresholds in unexposed **(C)** and NE mice **(D)** 2 weeks (2w) after NE. Error bars, mean ± SEM. Two-way ANOVA with a *post hoc* Tukey’s multiple comparisons test. Only significant two-way comparisons are labeled (*p* < 0.05). DTX-control: unexposed, *N* = 8; NE, *N* = 9. LOC-ablated: unexposed, *N* = 12; NE, *N* = 15. SAL-control: unexposed, *N* = 10; NE, *N* = 9.

**Figure S6:**
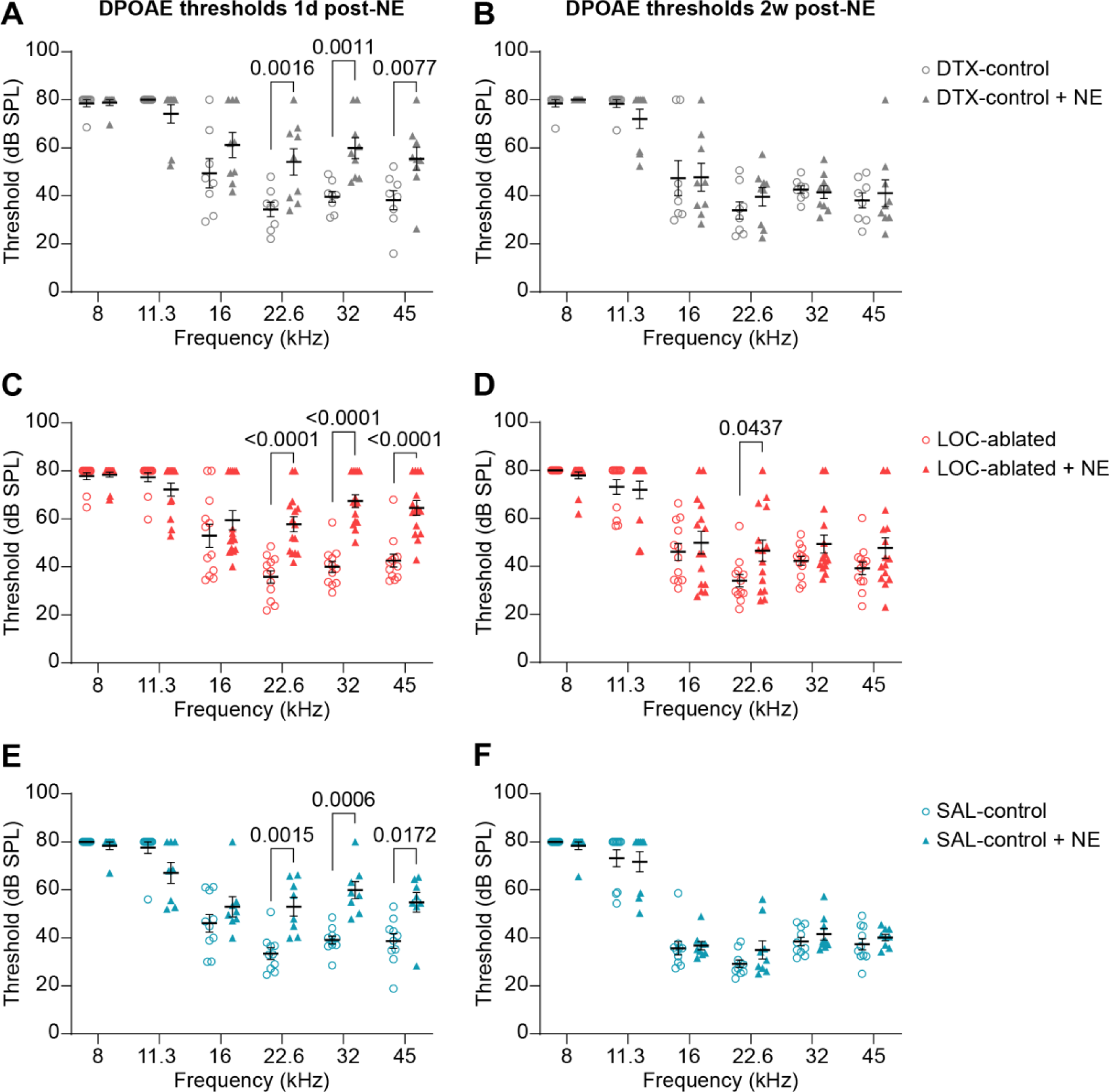
NE paradigm caused a temporary shift in DPOAE threshold. Comparisons of DPOAE thresholds between unexposed (open circles) and noise-exposed (NE, triangles) DTX-control **(A, B)**, LOC-ablated **(C, D)**, and SAL-control **(E, F)** mice 1 day (1d) **(A, C, E)** and 2 weeks (2w) **(B, D, F)** after NE. In all conditions, elevated DPOAE thresholds 1d after NE recovered 2w later, except at 22.6 kHz in LOC-ablated mice **(D)**. Error bars, mean ± SEM. Two-way ANOVA with Tukey’s *post hoc* multiple comparisons test. Only significant two-way comparisons are labeled (*p* < 0.05). Sample sizes are the same as **Fig. S5**.

## METHOD DETAILS

### Mice

All animal work was conducted in compliance with protocols approved by the Institutional Animal Care and Use Committee at Harvard Medical School. The following published mouse lines were used: *Rosa26^LSL-tdTomato^*(*Ai14*; Jax strain 007914) (92), *Rosa26^FSF-tdTomato^*(*Ai65F*; Jax strain 032864) (93), *Rosa26^RC::FLTG^*(Jax strain 026932) (39), *Rosa26^Sun1-GFP^*(Jax strain 021039) (94), and *Chat^CreΔNeo^*(Jax strain 031661) (95). Mice were maintained on a C57BL/J or mixed C57BL/J;129S6 background. *Mapt^ds-DTR^* mice (MGI:5911677) were obtained from the Ma Lab (96, 97). All mice used for LOC ablation experiments were heterozygous for the *Cdh23* mutation (*Cdh23^753G>A^*) associated with accelerated age-related hearing loss (98). Because they retain one unmutated copy of *Cdh23* (i.e., *Cdh23^753A/753G^*), our experimental mice are not expected to experience premature hearing loss. Regardless, all experiments were performed before animals reached 10w of age to ensure our results were not confounded by early-onset hearing loss.

Mice that received DTX or SAL were given 2 intraperitoneal injections 72h apart between 4–7w old. All mice in an experimental cohort received injections at the same time. DTX (Sigma D0564) was diluted in sterile saline to a concentration of 12.5 μg/mL and mice were given 50 μg/kg body weight. Mice given SAL received 4 μL/g body weight of sterile saline (comparable volume to the DTX solution administered).

### Generation of transgenic mouse lines

Both *Gata3^FlpO^* and *Ucn^Cre^* mouse lines were generated by homologous recombination via CRISPR/Cas9 targeting, using methods similar to those described previously (99, 100). *FlpO* (101) and *Cre* (102) sequences were directed to the endogenous locus of *Gata3* and *Ucn*, respectively, and inserted directly before the stop codon of each gene (**Fig. 2A, G**). Single-stranded homology repair templates (Genewiz) included sequences for a T2A peptide and SV40 nuclear localization sequence upstream of the *FlpO* or *Cre* recombinase sequence, as well as 400 bp homology arms on each side of the insertion construct. Each homology repair construct also induced a mutation at the targeted PAM site to prevent cutting of the repaired strand. In the *Ucn^Cre^*construct, we made a synonymous change in the coding sequence that did not alter the amino acid sequence of the peptide. For *Gata3^FlpO^*, we induced a C-to-G mutation within the gene’s 3’ UTR. Potential guide RNA sequences were selected based on their proximity to the construct insertion site and their bioinformatically predicted off-target effects (103). Guide RNAs were then screened *in vitro* to determine their efficacy at cutting a PCR-amplified fragment of target DNA. The final guides for targeting both *Ucn* (CCGCAUCAUAUUCGAUUCGG) and *Gata3* (UUGGAGACUCCUCACGCAUG) were synthesized as a combined CRISPR/tracr RNA molecule (Synthego). Finally, the guide RNA and homology repair templates were injected into zygotes of C57BL/6 mice along with Cas9 protein (IDT, Alt-R® S.p. Cas9 Nuclease V3; ca. 1081058) at Harvard’s Genome Modification Facility.

### Screening transgenic mouse lines

DNA was extracted from a tail clipping of each of the injected animals. We verified that each insertion was correctly targeted to the desired locus by PCR amplifying specific fragments of the construct and surrounding DNA. Next, we amplified and sequenced the entire inserted construct along with several hundred base pairs on either side of the insertion to verify that the homology repair construct was integrated into the locus as expected. Although the full constructs were inserted correctly in both *Gata3^FlpO^* and *Ucn^Cre^* mouse lines, we identified a few unexpected point mutations, likely caused by errors in the ssDNA synthesis process. In *Ucn^Cre^*: Ucn NM_001346010: c.312G>T (p.Glu104His); Cre: c.376G>T (p.Val126Phe); Cre: c.1029G>T (p.Gly343Gly). In *Gata3^FlpO^*: Gata3 NM_001417051: c.1375G>T (c.*43G>T in the Gata3 3’ UTR); FlpO: c.35C>A (p.Glu12Lys); Flpo:g.387G>A (p.Val129Val).

Upon verifying the insertion of *Gata3^FlpO^*, we genotyped subsequent generations with the following primer sequences: GAAGGCATCCAGACCCGAAA; CACGTCACCGCATGTTAGAAGA; AACGCAAGTAGAAGGGGTCG. Wild-type animals display a band at 336 bp. Animals homozygous for the *Gata3^FlpO^* allele yield a band at 291 bp. Heterozygous animals yield both the wild-type and mutant bands. We used the following primer sequences to genotype *Ucn^Cre^* animals: GGATCCGAATCTGCGATGGA;GCATCGACCGGTAATGCAGG;GGAGGCGAAATAGTCCCTC G. Animals homozygous for *Ucn^Cre^* display a band at 393 bp, whereas wild-type animals yield a band at 686 bp; heterozygotes show both the wild-type and mutant bands.

### Single-Nucleus Sequencing

#### Collection and Sequencing

Each round of single-nucleus sequencing included pooled tissue from 3–5 *Chat^CreΔNeo^;Rosa26^Sun1-GFP^* mice of both sexes between 6–10 weeks of age. In each case, tissue from noise-exposed animals and unexposed littermate controls was collected simultaneously and run on parallel lanes of the same 10x Genomics microfluidic chip. Two independent collections for the 1d post-NE timepoint included a total of 8 unexposed (5 male, 3 female) and 9 NE mice (6 male, 3 female). Four collections for the 7d post-NE timepoint included 14 unexposed (7 male, 7 female) and 15 NE mice (8 male, 7 female).

After being deeply anesthetized with isoflurane, mice were euthanized via rapid decapitation and brains were removed into ice-cold dissection buffer containing 0.25 M sucrose, 25 mM KCl, 5 mM MgCl_2_, 1M Tricine-KOH, 1 µM tetrodotoxin (TTX, Cayman Chemical 14964), 50 µM APV (Tocris 0106), and 20 µM DNQX (Sigma D0540). The ventral brainstem was dissected to include the entire SOC and nearby motor nuclei. Dissected brainstem tissue was pooled into a dounce homogenizer with a solution composed of 25 mM KCl, 5 mM MgCl2, 1M Tricine-KOH, 1 mM DTT (Sigma D0632), 150 µM spermine tetrahydrochloride (Sigma S1141), 500 µM spermidine trihydrochloride (Sigma S2501), 80 u/mL RNAsin Plus RNase Inhibitor (Promega N2615), and one tablet of protease inhibitor cocktail (Roche 11836170001). Partway through homogenization, IGEPAL CA-630 (Sigma I8896) was added to a final concentration of 0.32%. Manual homogenization continued until no chunks of tissue were visually apparent. The homogenized tissue was filtered through a 40 µm cell strainer and 5 mL of solution containing 50% iodixanol (OptiPrep Density Medium, Sigma D1556) with 7.5 mM KCl, 1.5 mM MgCl2, 6 mM Tricine-KOH, pH 7.8, and 80 u/mL RNAsin was added. The homogenate was then added to a density gradient of 30% and 40% iodixanol and spun for ∼25 minutes at 10,000 g at 4°C. Following the spin-down, the top layer was discarded and 400 µL was collected from the interface of the 30% and 40% iodixanol layers. Nuclei were diluted with 600 µL 1% BSA in PBS, stained with 10 µL of Draq7 (Abcam), and filtered through a 40 µm Flowmi cell strainer (Sigma-Aldrich). Nuclei were then sorted on a BD FACSAria II (BD Biosciences) by the Harvard Immunology Flow Cytometry Core to enrich for Sun1-GFP+ cholinergic nuclei. Sorted nuclei were centrifuged at 4°C for 5 min at 750 rpm and re-suspended in ∼40 µL of 1% BSA in PBS. Concentration of the sample was estimated by counting DAPI-labeled nuclei on a hemocytometer to determine appropriate cell-suspension volume for loading into a single-cell 3’ chip from 10x Genomics, following the manufacturer’s directions. For all collections, the maximum cell suspension volume was loaded. Nuclei were barcoded and processed using the Chromium Next GEM Single Cell 3’ Kit, v3.1 with G Chips and the Dual Index Kit TT, Set A (part numbers 1000268, 1000120, and 1000215). cDNA libraries were generated according to the 10x Genomics instructions. Libraries were pooled and sequenced on a NovaSeq S4 by the Harvard Bauer Core.

#### Alignment, filtering, and exclusion criteria

Sequencing reads were demultiplexed and aligned to a modified mm10-3.0.0 transcriptome that included intronic reads using the cellranger package from 10x Genomics (v. 7.0.0). Raw gene counts were then imported into the Seurat package (v. 4.2.1) in R (v.4.2.2). Cells and genes were initially filtered by eliminating all genes detected in fewer than 3 cells and removing cells with fewer than 500 unique genes. Because sequencing coverage varied somewhat between sequencing libraries, we used a standard-deviation based cutoff to remove potential multiplets: nuclei were excluded if the number of genes or unique molecular identifiers (UMI) detected was greater than two standard deviations above the mean. Following this filtering step, we eliminated additional low-quality cells by excluding nuclei with fewer than 1,000 UMI or 750 genes. Finally, we eliminated nuclei with a high fraction of genes mapping to the mitochondrial genome, as a high fraction of mitochondrial reads can indicate poor cell health. Because the fraction of nuclear reads mapping to the mitochondrial genome should be very low, we excluded all cells in which more than 1% of reads mapped to mitochondrial genes. At this point, we identified one collection round in which approximately 95% of nuclei from both control and experimental animals failed to pass our filtering criteria. Because the low number of cells passing QC indicates that these libraries were of low quality overall, the entire library was excluded from all analyses. Excluding the poor-quality collection round, approximately 8.6% of nuclei were eliminated by these filtering criteria, yielding a total of 99,519 single nuclei for downstream analysis.

#### Normalization and batch correction

Libraries were normalized independently using regularized negative binomial regression, implemented in the scTransform package in Seurat (v. 0.3.5) (104, 105). Normalized datasets were integrated with canonical correlation analysis (CCA) referenced to a subset of noise-exposed and control nuclei using 30 principal components (PCs) and 3,000 variable genes (106). After integrating the data, per-cell transcript counts were normalized by dividing the counts for each gene by the total counts for each cell. Next, these values were multiplied by 10,000 and natural-log transformed.

#### Clustering and differential expression analysis

Prior to clustering, we performed principal component analysis (PCA) on the integration vectors from the CCA pipeline above. We included 20 PCs as input to a graph-based clustering algorithm, implemented with the FindNeighbors and FindClusters algorithms in Seurat (106), using a resolution of 0.4. This analysis identified a single cluster each of MOC and LOC neurons. To refine our assessment of OCN subtypes, we subset this larger dataset to include only the 1,757 nuclei called as either MOCs or LOCs in the full dataset. We then repeated the PCA and clustering analysis on the subsetted data, again using 20 PCs and a resolution of 0.4.

Differentially expressed genes were identified with a nonparametric Wilcoxon rank-sum test on scTransform-normalized counts data followed by a Bonferroni post-hoc test for multiple comparisons, implemented with the FindMarkers function in Seurat.

### Noise Exposure

Noise exposure occurred in a custom plexiglass trapezoidal box located inside of a tabletop sound-attenuating chamber. These experiments occurred outside of the vivarium, requiring transport of mice to and from the noise exposure procedure room. Acoustic stimuli were digitally generated and amplified (Crown Xli 2500), then delivered from a speaker (JBL 2380A) positioned at the top of the plexiglass chamber. Each animal was placed in a wire mesh box on a mesh platform located in the center of the chamber, with up to four mice noise exposed in one session. Sound pressure level (SPL) was measured with a ¼” free-field microphone (PCB 378C01) that was calibrated prior to each exposure session (Larson-Davis CAL200). Mice were presented with an 8-16 kHz octave-band noise for 120 minutes. The mean intensities of the stimulus were as follows: sequencing experiments, 110.09 ± 0.47 dB SPL (absolute range); LOC ablation experiments, 97.09 ± 0.9 dB SPL (absolute range). Unexposed controls were left in their home cages in the vivarium.

### Auditory brainstem responses (ABR) and distortion product otoacoustic emissions (DPOAE)

All ABR and DPOAE measurements and analysis were conducted blind to the animal’s experimental condition. Both assays were conducted either one day and again two weeks after NE, or to littermate control animals that did not undergo the NE paradigm.

Mice were anesthetized with an intraperitoneal injection of ketamine (150 mg/kg body weight) and xylazine (10 mg/kg body weight) and placed on their side on a 37°C heating pad (ATC10000, World Precision Instruments) in a sound-attenuating, electrically shielded chamber for the duration of the experiment. Lubricant (GenTeal Tears, Alcon) was applied to both of the animal’s eyes to prevent drying. A dorsoventral incision at the intertragal notch of the pinna was used to expose the ear canal allowing for reliable probe placement. A 1/2–1/3 ketamine boost of the initial induction volume was used to maintain anesthesia at 30-minute intervals or at the first sign of movement (e.g. whisking or twitching).

Acoustic stimuli were delivered and measured using a custom acoustic system consisting of a microphone and dual earphone assembly coupled to a probe tube. The probe end of the acoustic system was placed directly over the ear canal. Stimuli were digitally generated and then amplified by an SA1 stereo power amp (Tucker Davis Technologies) to drive the acoustic system. The probe tube microphone was amplified by an MA3 microphone amp (Tucker Davis Technologies) and sent to an I/O board installed into a PXI chassis (National Instruments). The custom acoustic system and LabVIEW software (EPL Cochlear Function Test Suite and ABR Peak Analysis Software) were provided by the Engineering Core of the Eaton-Peabody Laboratories at Massachusetts Eye and Ear Infirmary (MEEI), Boston.

ABRs were recorded after implanting subdermal electrodes into the animal: an active electrode at the vertex, a reference electrode near the pinna, and a ground electrode near the tail. Bioelectrical signals were amplified through an AC pre-amplifier (Grass Technologies) before digitization. Acoustic stimuli consisted of 5 ms tone pips with a repetition rate of 30 s^-1^ with a 0.5 ms rise-fall time at the following frequencies (in kHz): 8, 11.3, 16, 22.6, 32, and 45. Stimuli were presented in alternating polarity to cancel out the cochlear microphonic potential. Cochlear Function Test Suite (CFTS) software (version 2.36, MEEI, Boston) amplified (x104), filtered (0.3– 3 kHz passband), and averaged 512 responses at each SPL.

DPOAEs were recorded in response to primary tones f1 and f2 (frequency ratio f2/f1 = 1.2, SPL level of L1 = L2 + 10 dB), and f2 consisted of 5.6, 8, 11.3, 16, 22.6, and 32 kHz tones that were incremented in 10 dB steps from 10 to 70 dB SPL. The cubic distortion product 2f1-f2 was extracted by Fourier analysis of the ear-canal sound pressure after waveform and spectral averaging. DPOAE threshold was defined as the f1 level required to produce a DPOAE of 5 dB SPL.

### Immunohistochemistry

#### Brainstem histology

Mice were deeply anesthetized with isoflurane prior to transcardial perfusion with 5–15ml of ice-cold 4% PFA in PBS. Brains were removed and post-fixed overnight at 4°C, then rinsed 3X with 1X PBS.

For histology on *Gata3^FlpO^* and *Ucn^Cre^* tissue (**Fig. 2**), brains were cryoprotected through a sucrose gradient of 10%, 20%, then 30% sucrose in 1X PBS, followed by a 30% sucrose/NEG mix, before embedding in 100% NEG and freezing. Tissue was cryosectioned at 20–25 μm on a Leica CM3050S cryostat. Sections were permeabilized and blocked for 1–2 h at room temperature (RT) in 1X PBS with 0.5% Triton-X (0.5% PBST) and 5–10% normal donkey serum (NDS, Jackson ImmunoResearch 017-000-121). Primary and secondary antibodies were diluted in 0.5% PBST and 5% NDS. Tissue was incubated in primary antibodies overnight at 4°C, followed by 3 rinses with 1X PBS, and incubation with secondary antibodies for 1.5–3 h at RT. Primary and secondary antibodies were mixed in 0.5% PBST with 5% NDS. Following secondary antibody incubation, slides were incubated with DAPI diluted 1:10,000 in 1X PBS for 10 minutes at RT.

For histology on *Gata3∩Ucn* tissue (**Figs. 2L, M & 3B**), brains were embedded in a bovine serum albumin (BSA) gelatin embedding media (0.4% gelatin, 23% BSA, 26% formalin, 1.4% glutaraldehyde) and sectioned 60 μm thick on a Leica VT1000S vibratome. Free-floating sections were permeabilized and blocked in 5% NDS in 0.2% PBST for 1h at RT before incubating in primary antibodies overnight at 4°C. Following primary antibody incubation, sections were washed 3X with 1X PBS, then incubated with secondary antibodies for 1.5–3 h at RT. Tissue was then rinsed 3X with 1X PBS and mounted using Fluoromount-G (Southern Biotech 0100-01). Primary and secondary antibodies were mixed in 5% NDS in 0.2% PBST. Primary antibodies used were Goat anti-ChAT (Millipore AB144P, 1:200), Rabbit anti-dsRed (Takara 632496, 1:1,000–1:2,000), and Chicken anti-GFP (Aves GFP-1020, 1:2,000). Secondary antibodies used were Donkey anti-Goat 488 (Jackson ImmunoResearch 705-545-147, 1:800), Donkey anti-Rabbit 568 (Abcam ab175470 or ThermoFisher A31573, 1:800–1:1,000), and Donkey anti-Chicken 488 (Jackson ImmunoResearch 703-545-155, 1:800).

#### Cochlea histology

Mice were deeply anesthetized with isoflurane prior to transcardial perfusion with 5–15 ml of ice-cold 4% PFA in PBS. Temporal bones were removed, post-fixed overnight at 4°C, then rinsed 3X with 1X PBS and decalcified in 120 mM EDTA (Ethylenediaminetetraacetic acid, Avantar E177) for 48–72 h at 4°C. Following decalcification, temporal bones were rinsed with 1X PBS and dissected into three cochlear turns. For characterization of transgenic mouse lines in **Fig. 2**, dissected cochlea were permeabilized by treatment with protease for 30 min at 37°C (ACD RNAscope Protease III; ca.322340). All cochlear turns, regardless of protease treatment, were cryoprotected in 30% sucrose for at least 20 min at RT, then permeabilized by flash-freezing on dry ice. After thawing in a RT water bath, tissue was rinsed 3X in 1X PBS and permeabilized and blocked in 5% NDS in 0.3–1% PBST for 45–120 min at RT. For cochlea labeled for presynaptic ribbons (Ctbp2, **Fig. 6**), tissue was then rinsed 3X in 0.02% PBST, incubated with Donkey anti-Mouse IgG Fab Fragments (1:25; Jackson ImmunoResearch 715-007-003) in 0.3% PBST for 45– 90 minutes at RT, then rinsed again 3X with 0.02% PBST at RT. Otherwise, tissue proceeded directly from blocking solution to primary antibody solution. Tissue was incubated with primary antibodies at 37°C overnight in 1% NDS and 0.3–0.5% PBST, then rinsed 3X in 0.5–1% PBST before incubating with secondary antibodies for 2h overnight at 37°C. Tissue was rinsed 2–3X with 0.5–1% PBST, then up to 2X with 1X PBS before mounting in Fluoromount-G in wells cut into 0.81 mm-thick vinyl tape (3M, catolog number 471). Primary antibodies used: Guinea pig anti-Parvalbumin (Synaptic Systems 195-004, 1:1000), Chicken anti-GFP (Aves GFP-1020, 1:2,000); Mouse IgG1 anti-Ctbp2 (BD Biosciences 612044, 1:500); Goat Anti-Chat (EMD Millipore AB144P, 1:1000); Rabbit Anti-dsRed (Takara 632496, 1:1000); Goat Anti-Calb2 (Swant CG1, 1:1000). Secondary antibodies used: Donkey anti-Guinea Pig Alexa 405 (Sigma SAB4600230-50, 1:1000); Donkey anti-Chicken Alexa 488 (Jackson ImmunoResearch 703-545-155, 1:1000); Donkey anti-Mouse Alexa 647 (Thermo Fisher A31571, 1:1000); Donkey anti-Rabbit Alexa 568 (Abcam ab175470 or ThermoFisher A31573, 1:800-1:1,000); Donkey anti-Mouse Alexa 647 (Thermo Fisher A31571, 1:1000); Donkey anti-Goat Alexa 488 (Jackson ImmunoResearch 705-545-147, 1:1000).

### Imaging

Brainstem sections were imaged on an Olympus VS120 slide scanner using the UPLSAPO 10x/0.40 objective or on a Zeiss LSM800 confocal microscope Plan-Apochromatic 20x/0.8 M27 objective. Cochlea were imaged on a Leica SP8 confocal microscope using an HC PL APO CS2 10x/0.4 objective or an HC PL APO CS2 40x/1.3 oil objective; or on a Zeiss LSM800 confocal microscope using an EC-Plan NeoFluar 10x/0.30 M27 objective and a PlanApo 63x/1.4 oil objective. Images were processed in the Fiji distribution (107) of ImageJ or Zeiss Zen (v2.3 or 3.8) software.

### Ribbon Quantification

To analyze putative SGN synapses in mice for the LOC ablation experiments, animals were collected 1–5 d following the second ABR and cochlear tissue was processed as described above. To determine frequency regions along the length of the cochlea, low-magnification images of each cochlear turn were acquired on a Zeiss LSM800 confocal microscope using a EC-Plan NeuroFluar 10x/0.30 M27 objective. Images were imported into Fiji (107) using the Bio-Formats plugin (108) and a cochlear frequency map was generated using the measure_line plugin distributed by Eaton Peabody Labs, MEEI. High-magnification images were then acquired at 8, 16, and 45.2 kHz frequency regions using PlanApo 63x/1.4 oil objective on a Zeiss LSM800 confocal, and imported into Imaris (Bitplane AG, v. 9.9.0). Each image contained a ∼100μm length of cochlea centered at the respective frequency. Ctbp2 puncta in unexposed and NE LOC-ablated and DTX-control mice were semi-automatically identified using the spots function with local contrast background subtraction and an estimated spot diameter of 0.495 μm. Spots were manually reviewed by a researcher blind to condition and corrected as needed. Coordinates were created using the reference frame function for each IHC, with 0, 0, 0 at the base of the hair cell, with the XZ plane bisecting the hair cell through its plane of symmetry along the modiolar-pillar axis, a YZ plane bisecting the nucleus along the basilar-cuticular axis of the IHC, and an XY plane tangential to the basolateral pole of the hair cell running the length of the cochlea. This allowed for comparison of the modiolar-pillar distribution of Ctpb2 puncta (no substantive differences were found between conditions). Ctbp2 spots were assigned by the researcher to their respective IHC and the hair-cell-centric reference frames were used to assess the spatial distribution of Ctbp2 spots within a given IHC. Dying IHCs were removed from analysis based on the following criteria: qualitatively smaller than average IHCs, displaced towards the cuticular plate, and containing three or fewer Ctbp2 puncta that were all tightly clustered to the basal pole of the IHC. Data were exported from Imaris into .csv files and python (v 3.10) with pandas (v 2.0.3) was used to filter relevant metrics into a new .csv. Statistical analysis was then performed and graphical plots produced using GraphPad Prism (v 10.1.1).

## RESOURCE TABLE

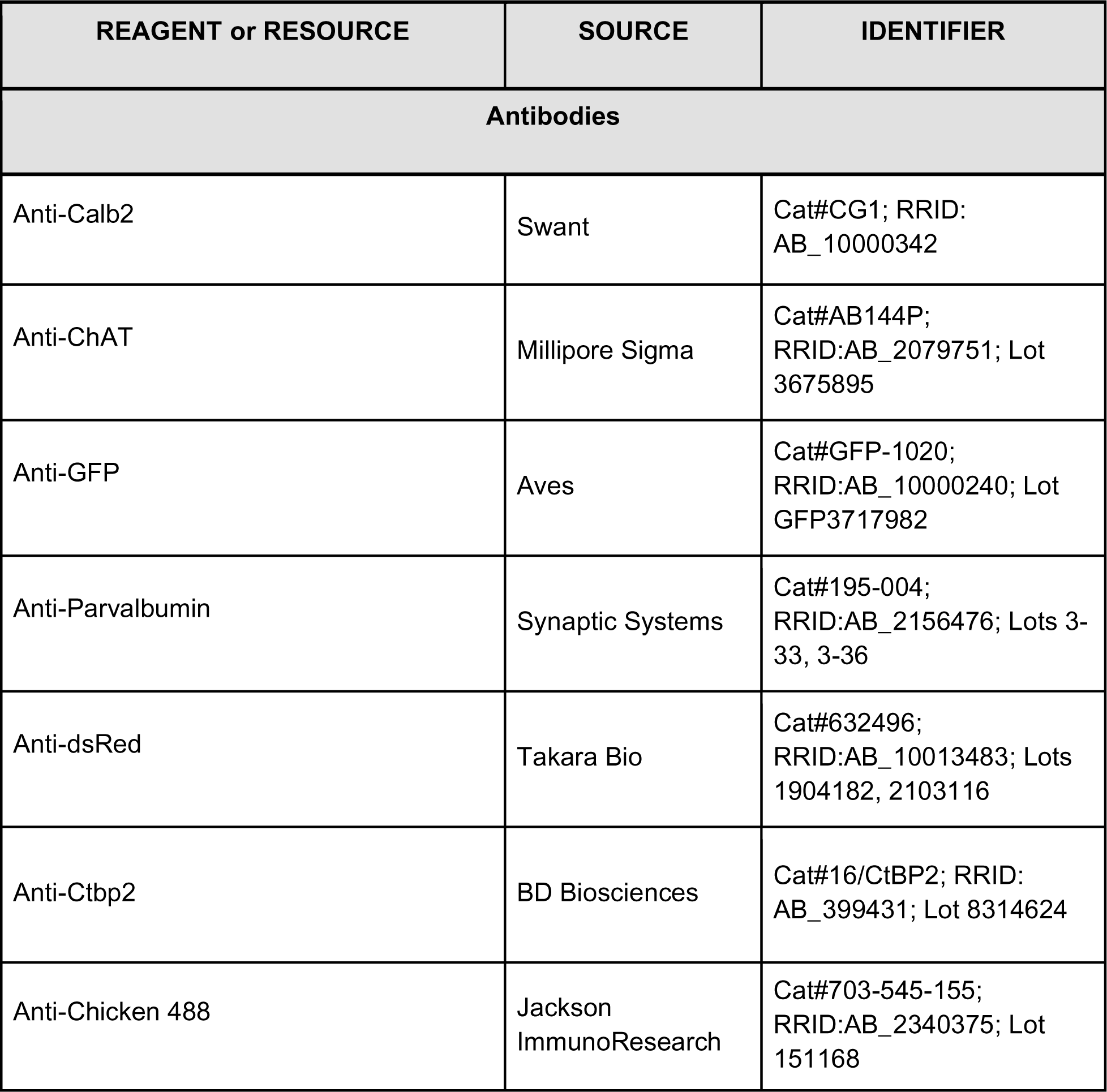
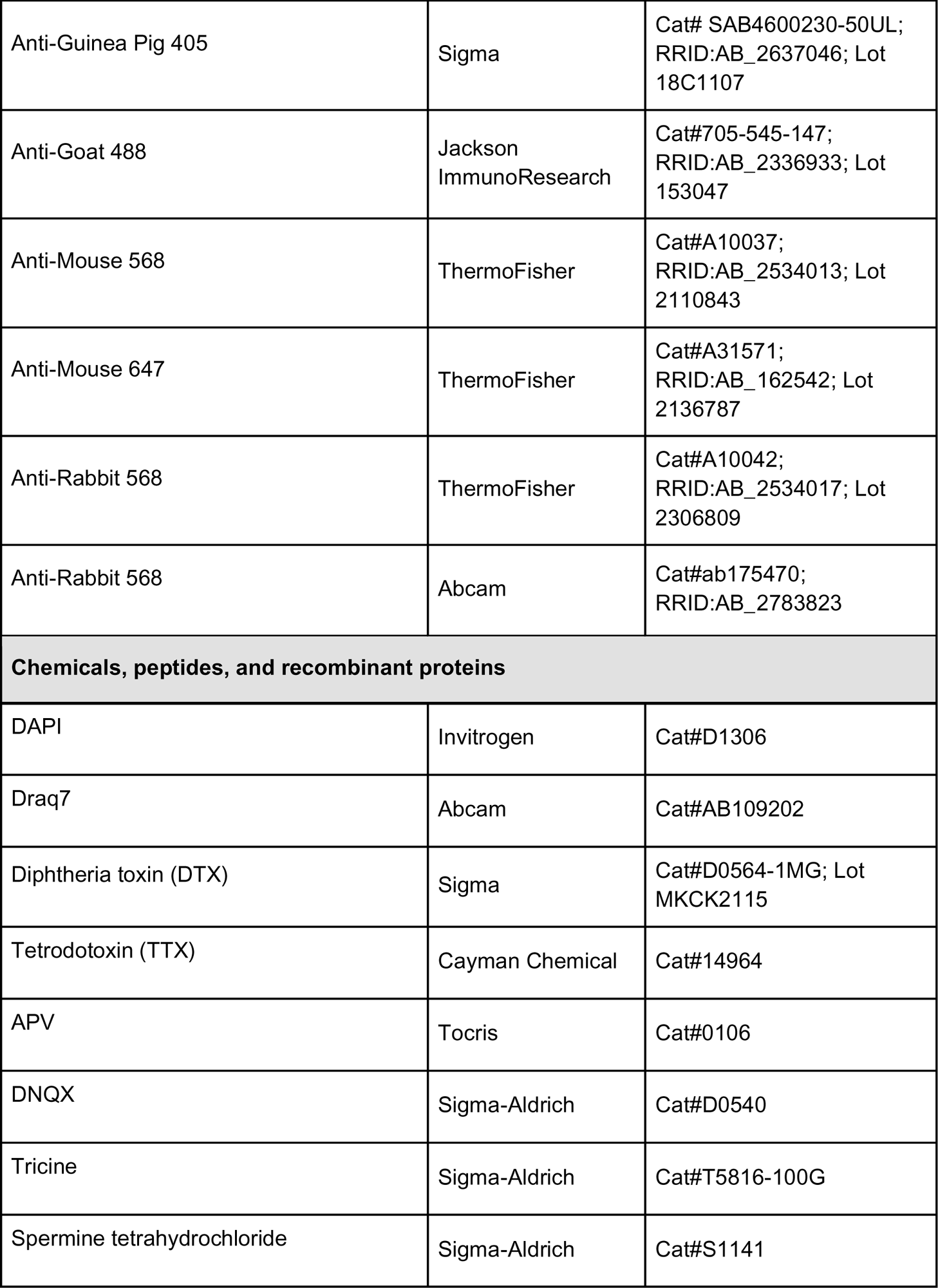
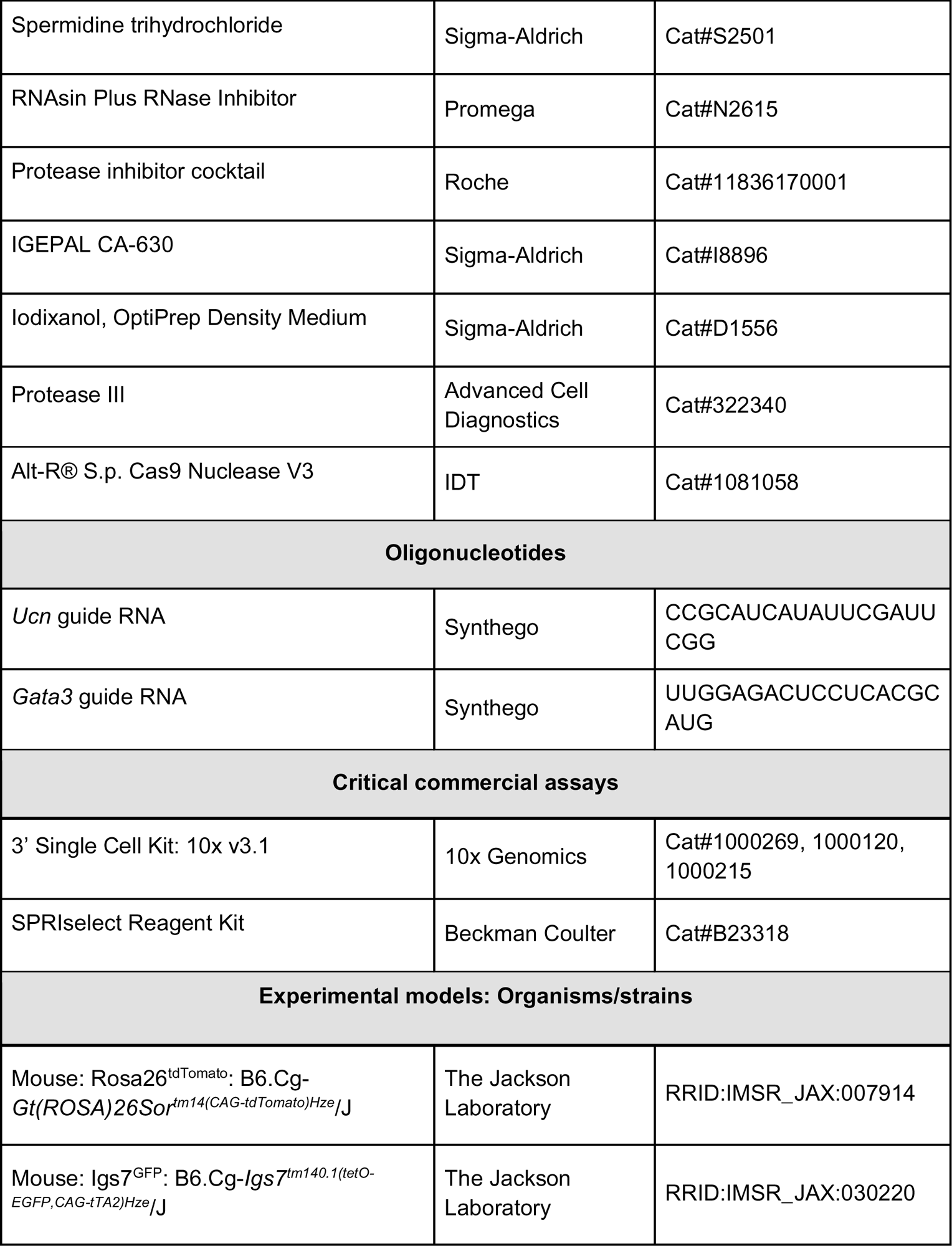
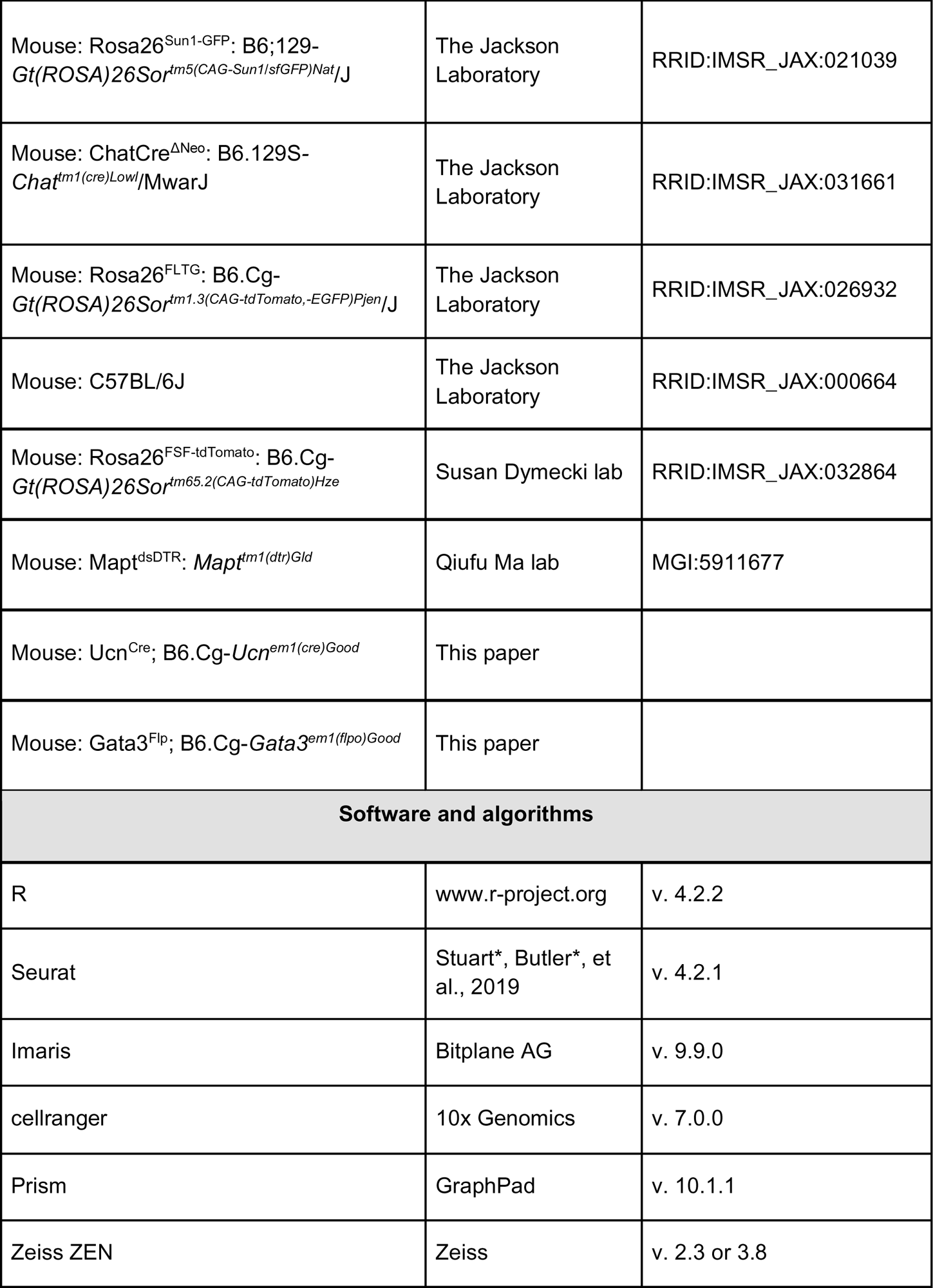
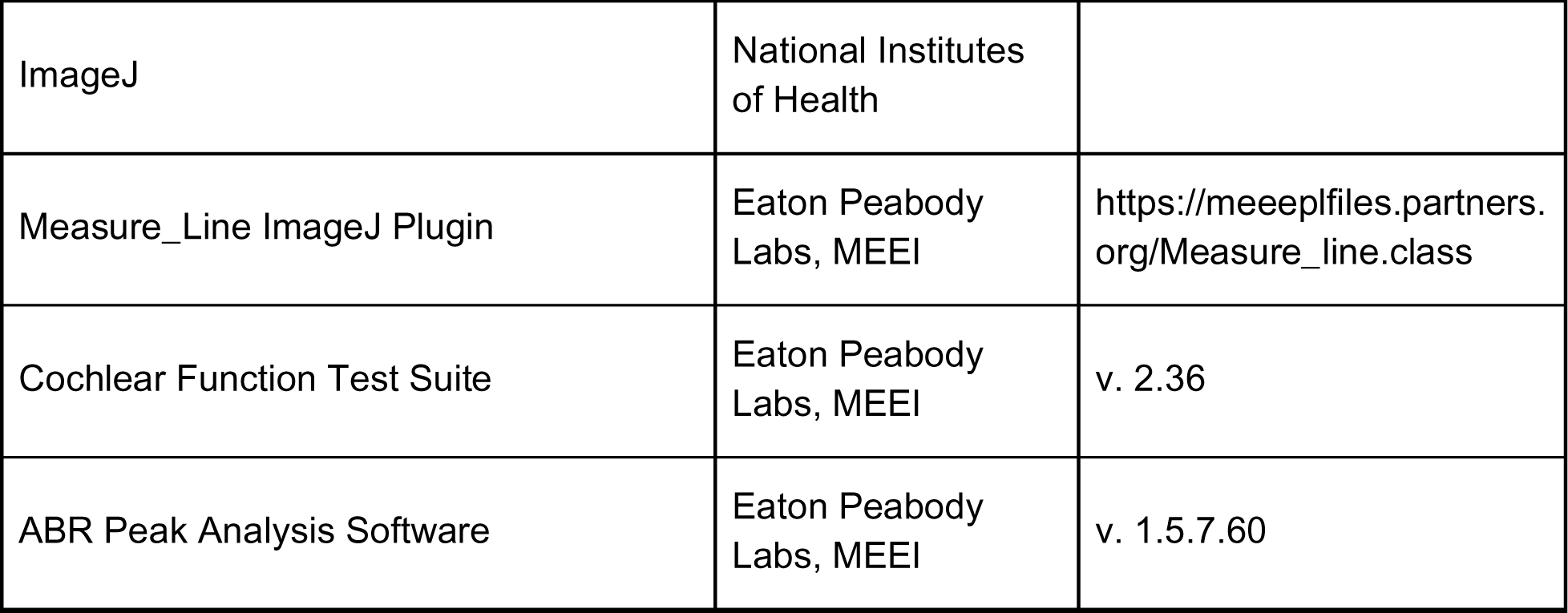

## Notes

### Competing Interest Statement

The authors have declared no competing interest.

## REFERENCES

1. J. J. Guinan, Olivocochlear efferents: Their action, effects, measurement and uses, and the impact of the new conception of cochlear mechanical responses. Hear. Res. 362, 38–47 (2018).

2. A. M. Lauer, S. V. Jimenez, P. H. Delano, Olivocochlear efferent effects on perception and behavior. Hear. Res. 419, 108207 (2022).

3. G. E. Romero, L. O. Trussell, Central circuitry and function of the cochlear efferent systems. Hear. Res. 425, 108516 (2022).

4. M. C. Brown, Morphology of labeled efferent fibers in the guinea pig cochlea. J. Comp. Neurol. 260, 605–618 (1987).

5. M. M. Frank, et al., Experience-dependent flexibility in a molecularly diverse central-to-peripheral auditory feedback system. eLife 12, e83855 (2023).

6. J. J. Guinan, W. B. Warr, B. E. Norris, Differential olivocochlear projections from lateral versus medial zones of the superior olivary complex. J. Comp. Neurol. 221, 358–370 (1983).

7. Y. Hua, et al., Electron Microscopic Reconstruction of Neural Circuitry in the Cochlea. Cell Rep. 34, 108551 (2021).

8. W. B. Warr, J. J. Guinan, Efferent innervation of the organ of corti: two separate systems. Brain Res. 173, 152–155 (1979).

9. M. L. Gifford, J. J. Guinan, Effects of electrical stimulation of medial olivocochlear neurons on ipsilateral and contralateral cochlear responses. Hear. Res. 29, 179– 194 (1987).

10. D. C. Mountain, Changes in endolymphatic potential and crossed olivocochlear bundle stimulation alter cochlear mechanics. Science 210, 71–72 (1980).

11. R. D. Rabbitt, W. E. Brownell, Efferent modulation of hair cell function. Curr. Opin. Otolaryngol. Head Neck Surg. 19, 376–381 (2011).

12. J. H. Siegel, D. O. Kim, Efferent neural control of cochlear mechanics? Olivocochlear bundle stimulation affects cochlear biomechanical nonlinearity. Hear. Res. 6, 171–182 (1982).

13. J. Taranda, et al., A point mutation in the hair cell nicotinic cholinergic receptor prolongs cochlear inhibition and enhances noise protection. PLoS Biol. 7, e18 (2009).

14. J. A. Groff, M. C. Liberman, Modulation of Cochlear Afferent Response by the Lateral Olivocochlear System: Activation Via Electrical Stimulation of the Inferior Colliculus. J. Neurophysiol. 90, 3178–3200 (2003).

15. J. S. Wu, et al., Sound exposure dynamically induces dopamine synthesis in cholinergic LOC efferents for feedback to auditory nerve fibers. eLife 9, 1–27 (2020).

16. T. T. Hickman, K. Hashimoto, L. D. Liberman, M. C. Liberman, Synaptic migration and reorganization after noise exposure suggests regeneration in a mature mammalian cochlea. Sci. Rep. 10, 19945 (2020).

17. S. G. Kujawa, M. C. Liberman, Translating animal models to human therapeutics in noise-induced and age-related hearing loss. Hear. Res. 377, 44–52 (2019).

18. S. G. Kujawa, M. C. Liberman, Adding insult to injury: cochlear nerve degeneration after “temporary” noise-induced hearing loss. J. Neurosci. Off. J. Soc. Neurosci. 29, 14077–14085 (2009).

19. M. C. Liberman, L. W. Dodds, Single-neuron labeling and chronic cochlear pathology. III. Stereocilia damage and alterations of threshold tuning curves. Hear. Res. 16, 55–74 (1984).

20. L. Shi, et al., Ribbon synapse plasticity in the cochleae of Guinea pigs after noise-induced silent damage. PloS One 8, e81566 (2013).

21. Y. Wang, K. Hirose, M. C. Liberman, Dynamics of noise-induced cellular injury and repair in the mouse cochlea. JARO - J. Assoc. Res. Otolaryngol. 3, 248–268 (2002).

22. M. C. Liberman, S. G. Kujawa, Cochlear synaptopathy in acquired sensorineural hearing loss: Manifestations and mechanisms. Hear. Res. 349, 138–147 (2017).

23. L. E. Boero, et al., Enhancement of the Medial Olivocochlear System Prevents Hidden Hearing Loss. J. Neurosci. 38, 7440–7451 (2018).

24. A. Fuente, The olivocochlear system and protection from acoustic trauma: A mini literature review. Front. Syst. Neurosci. 9, 1–6 (2015).

25. S. F. Maison, H. Usubuchi, M. C. Liberman, Efferent Feedback Minimizes Cochlear Neuropathy from Moderate Noise Exposure. J. Neurosci. 33, 5542–5552 (2013).

26. X. Y. Zheng, D. Henderson, S. L. McFadden, B. H. Hu, The role of the cochlear efferent system in acquired resistance to noise-induced hearing loss. Hear. Res. 104, 191–203 (1997).

27. K. N. Darrow, S. F. Maison, M. C. Liberman, Selective Removal of Lateral Olivocochlear Efferents Increases Vulnerability to Acute Acoustic Injury. J. Neurophysiol. 97, 1775–1785 (2007).

28. S. G. Kujawa, M. C. Liberman, Conditioning-Related Protection From Acoustic Injury: Effects of Chronic Deefferentation and Sham Surgery. J. Neurophysiol. 78, 3095–3106 (1997).

29. T. Arnold, E. Oestreicher, K. Ehrenberger, D. Felix, GABA(A) receptor modulates the activity of inner hair cell afferents in guinea pig cochlea. Hear. Res. 125, 147– 153 (1998).

30. G. P. Bailey, W. F. Sewell, Calcitonin Gene-Related Peptide Suppresses Hair Cell Responses to Mechanical Stimulation in the *Xenopus* Lateral Line Organ. J. Neurosci. 20, 5163–5169 (2000).

31. J. Ruel, et al., Dopamine inhibition of auditory nerve activity in the adult mammalian cochlea. Eur. J. Neurosci. 14, 977–986 (2001).

32. W. F. Sewell, “Pharmacology and Neurochemistry of Olivocochlear Efferents” in Auditory and Vestibular Efferents, D. K. Ryugo, R. R. Fay, A. N. Popper, Eds. (Springer, 2011), pp. 83–101.

33. X. Niu, B. Canlon, Activation of tyrosine hydroxylase in the lateral efferent terminals by sound conditioning. Hear. Res. 174, 124–132 (2002).

34. K. A. Fernandez, P. W. C. Jeffers, K. Lall, M. C. Liberman, S. G. Kujawa, Aging after Noise Exposure: Acceleration of Cochlear Synaptopathy in “Recovered” Ears. J. Neurosci. 35, 7509–7520 (2015).

35. A. Kaiser, O. Alexandrova, B. Grothe, Urocortin-expressing olivocochlear neurons exhibit tonotopic and developmental changes in the auditory brainstem and in the innervation of the cochlea. J. Comp. Neurol. 519, 2758–2778 (2011).

36. J. S. Duncan, K.-C. Lim, J. D. Engel, B. Fritzsch, Limited inner ear morphogenesis and neurosensory development are possible in the absence of GATA3. Int. J. Dev. Biol. 55, 297–303 (2011).

37. J. S. Duncan, B. Fritzsch, Continued expression of GATA3 is necessary for cochlear neurosensory development. PloS One 8, e62046 (2013).

38. A. Karis, et al., Transcription factor GATA-3 alters pathway selection of olivocochlear neurons and affects morphogenesis of the ear. J. Comp. Neurol. 429, 615–630 (2001).

39. N. W. Plummer, et al., Expanding the power of recombinase-based labeling to uncover cellular diversity. Dev. Camb. Engl. 142, 4385–4393 (2015).

40. H. Konishi, et al., Exposure to diphtheria toxin during the juvenile period impairs both inner and outer hair cells in C57BL/6 mice. Neuroscience 351, 15–23 (2017).

41. Y. Song, et al., Activity-dependent regulation of prestin expression in mouse outer hair cells. J. Neurophysiol. 113, 3531–3542 (2015).

42. A. Vavakou, N. P. Cooper, M. van der Heijden, The frequency limit of outer hair cell motility measured in vivo. eLife 8, e47667 (2019).

43. J. L. Puel, J. Ruel, C. Gervais d’Aldin, R. Pujol, Excitotoxicity and repair of cochlear synapses after noise-trauma induced hearing loss. Neuroreport 9, 2109–2114 (1998).

44. R. Pujol, J. L. Puel, C. Gervais d’Aldin, M. Eybalin, Pathophysiology of the glutamatergic synapses in the cochlea. Acta Otolaryngol. (Stockh.) 113, 330–334 (1993).

45. R. Pujol, J. L. Puel, Excitotoxicity, synaptic repair, and functional recovery in the mammalian cochlea: a review of recent findings. Ann. N. Y. Acad. Sci. 884, 249– 254 (1999).

46. L. D. Liberman, J. Suzuki, M. C. Liberman, Dynamics of cochlear synaptopathy after acoustic overexposure. J. Assoc. Res. Otolaryngol. JARO 16, 205–219 (2015).

47. S. Maison, M. C. Liberman, Predicting vulnerability to acoustic injury with a noninvasive assay of olivocochlear reflex strength. J. Neurosci. Off. J. Soc. Neurosci. 20, 4701–7 (2000).

48. S. F. Maison, A. E. Luebke, M. C. Liberman, J. Zuo, Efferent protection from acoustic injury is mediated via alpha9 nicotinic acetylcholine receptors on outer hair cells. J. Neurosci. Off. J. Soc. Neurosci. 22, 10838–10846 (2002).

49. R. Rajan, Involvement of cochlear efferent pathways in protective effects elicited with binaural loud sound exposure in cats. J. Neurophysiol. 74, 582–597 (1995).

50. R. Rajan, Effect of electrical stimulation of the crossed olivocochlear bundle on temporary threshold shifts in auditory sensitivity. II. Dependence on the level of temporary threshold shifts. J. Neurophysiol. 60, 569–579 (1988).

51. S. F. Maison, et al., Overexpression of SK2 channels enhances efferent suppression of cochlear responses without enhancing noise resistance. J. Neurophysiol. 97, 2930–2936 (2007).

52. H.-B. Zhao, et al., Efferent neurons control hearing sensitivity and protect hearing from noise through the regulation of gap junctions between cochlear supporting cells. J. Neurophysiol. 127, 313–327 (2022).

53. M. C. Liberman, The olivocochlear efferent bundle and susceptibility of the inner ear to acoustic injury. J. Neurophysiol. 65, 123–132 (1991).

54. M. C. Liberman, Effects of chronic cochlear de-efferentation on auditory-nerve response. Hear. Res. 49, 209–223 (1990).

55. M. C. Brown, “Anatomy of olivocochlear neurons” in Auditory and Vestibular Efferents, Springer Handbook of Auditory Research., D. K. Ryugo, R. R. Fay, Eds. (Springer New York, 2011), pp. 17–37.

56. K. Suthakar, M. C. Liberman, Auditory-nerve responses in mice with noise-induced cochlear synaptopathy. J. Neurophysiol. 126, 2027–2038 (2021).

57. R. Nouvian, M. Eybalin, J.-L. Puel, Cochlear efferents in developing adult and pathological conditions. Cell Tissue Res. 361, 301–309 (2015).

58. N. Dedic, A. Chen, J. M. Deussing, The CRF Family of Neuropeptides and their Receptors - Mediators of the Central Stress Response. Curr. Mol. Pharmacol. 11, 4–31 (2018).

59. V. Kormos, B. Gaszner, Role of neuropeptides in anxiety, stress, and depression: from animals to humans. Neuropeptides 47, 401–419 (2013).

60. F. Reichmann, P. Holzer, Neuropeptide Y: A stressful review. Neuropeptides 55, 99–109 (2016).

61. S. Udit, K. Blake, I. M. Chiu, Somatosensory and autonomic neuronal regulation of the immune response. Nat. Rev. Neurosci. 23, 157–171 (2022).

62. Y. Wang, M. C. Liberman, Restraint stress and protection from acoustic injury in mice. Hear. Res. 165, 96–102 (2002).

63. A. R. Dörrbaum, L. Kochen, J. D. Langer, E. M. Schuman, Local and global influences on protein turnover in neurons and glia. eLife 7, e34202 (2018).

64. T. J. Eisen, et al., The Dynamics of Cytoplasmic mRNA Metabolism. Mol. Cell 77, 786–799.e10 (2020).

65. N. L. Jorstad, et al., Transcriptomic cytoarchitecture reveals principles of human neocortex organization. Science 382, eadf6812 (2023).

66. G. Stanley, O. Gokce, R. C. Malenka, T. C. Südhof, S. R. Quake, Continuous and Discrete Neuron Types of the Adult Murine Striatum. Neuron 105, 688–699.e8 (2020).

67. F. Xie, S. Jain, S. Butrus, K. Shekhar, S. L. Zipursky, Vision sculpts a continuum of L2/3 cell types in the visual cortex during the critical period. bioRxiv, 2023.12.18.572244 (2023).

68. M. Zhang, et al., Spatially resolved cell atlas of the mouse primary motor cortex by MERFISH. Nature 598, 137–143 (2021).

69. G. P. Bailey, W. F. Sewell, Pharmacological Characterization of the CGRP Receptor in the Lateral Line Organ of Xenopus laevis. J. Assoc. Res. Otolaryngol. 1, 82–88 (2000).

70. D. Felix, K. Ehrenberger, The efferent modulation of mammalian inner hair cell afferents. Hear. Res. 64, 1–5 (1992).

71. K. Ito, D. Dulon, Nonselective cation conductance activated by muscarinic and purinergic receptors in rat spiral ganglion neurons. Am. J. Physiol. Cell Physiol. 282, C1121–1135 (2002).

72. C. G. Le Prell, L. F. Hughes, D. F. Dolan, S. C. Bledsoe, Effects of Calcitonin-Gene-Related-Peptide on Auditory Nerve Activity. Front. Cell Dev. Biol. 9, 752963 (2021).

73. S. F. Maison, et al., Muscarinic signaling in the cochlea: presynaptic and postsynaptic effects on efferent feedback and afferent excitability. J. Neurosci. Off. J. Soc. Neurosci. 30, 6751–6762 (2010).

74. S. F. Maison, R. B. Emeson, J. C. Adams, A. E. Luebke, M. C. Liberman, Loss of αCGRP Reduces Sound-Evoked Activity in the Cochlear Nerve. J. Neurophysiol. 90, 2941–2949 (2003).

75. W. F. Sewell, P. A. Starr, Effects of calcitonin gene-related peptide and efferent nerve stimulation on afferent transmission in the lateral line organ. J. Neurophysiol. 65, 1158–1169 (1991).

76. S. F. Maison, et al., Dopaminergic Signaling in the Cochlea: Receptor Expression Patterns and Deletion Phenotypes. J. Neurosci. 32, 344–355 (2012).

77. C. Petitpré, et al., Neuronal heterogeneity and stereotyped connectivity in the auditory afferent system. Nat. Commun. 9, 3691 (2018).

78. B. R. Shrestha, et al., Sensory Neuron Diversity in the Inner Ear Is Shaped by Activity. Cell 174, 1229–1246.e17 (2018).

79. S. Sun, et al., Hair Cell Mechanotransduction Regulates Spontaneous Activity and Spiral Ganglion Subtype Specification in the Auditory System. Cell 174, 1247–1263.e15 (2018).

80. C. E. Graham, J. Basappa, D. E. Vetter, A corticotropin-releasing factor system expressed in the cochlea modulates hearing sensitivity and protects against noise-induced hearing loss. Neurobiol. Dis. 38, 246–258 (2010).

81. C. E. Graham, D. E. Vetter, The Mouse Cochlea Expresses a Local Hypothalamic-Pituitary-Adrenal Equivalent Signaling System and Requires Corticotropin-Releasing Factor Receptor 1 to Establish Normal Hair Cell Innervation and Cochlear Sensitivity. J. Neurosci. 31, 1267–1278 (2011).

82. H. Liu, et al., Cell-Specific Transcriptome Analysis Shows That Adult Pillar and Deiters’ Cells Express Genes Encoding Machinery for Specializations of Cochlear Hair Cells. Front. Mol. Neurosci. 11 (2018).

83. D. E. Vetter, et al., Urocortin-deficient mice show hearing impairment and increased anxiety-like behavior. Nat. Genet. 31, 363–369 (2002).

84. C. E. Graham, J. Basappa, S. Turcan, D. E. Vetter, The Cochlear CRF Signaling Systems and their Mechanisms of Action in Modulating Cochlear Sensitivity and Protection Against Trauma. Mol. Neurobiol., 1–24 (2011).

85. S. R. Kitcher, A. M. Pederson, C. J. C. Weisz, Diverse identities and sites of action of cochlear neurotransmitters. Hear. Res. 419, 108278 (2022).

86. B. Milon, et al., A cell-type-specific atlas of the inner ear transcriptional response to acoustic trauma. Cell Rep. 36, 109758 (2021).

87. V. Rai, et al., The immune response after noise damage in the cochlea is characterized by a heterogeneous mix of adaptive and innate immune cells. Sci. Rep. 10, 15167 (2020).

88. L. Souza-Moreira, J. Campos-Salinas, M. Caro, E. Gonzalez-Rey, Neuropeptides as Pleiotropic Modulators of the Immune Response. Neuroendocrinology 94, 89– 100 (2011).

89. A. N. van den Pol, Neuropeptide transmission in brain circuits. Neuron 76, 98–115 (2012).

90. H. Xiong, et al., Probing Neuropeptide Volume Transmission In Vivo by Simultaneous Near-Infrared Light-Triggered Release and Optical Sensing. Angew. Chem. Int. Ed. 61, e202206122 (2022).

91. S. X. Zhang, et al., Competition between stochastic neuropeptide signals calibrates the rate of satiation. bioRxiv, 2023.07.11.548551 (2023).

92. L. Madisen, et al., A robust and high-throughput Cre reporting and characterization system for the whole mouse brain. Nat. Neurosci. 13, 133–140 (2010).

93. T. L. Daigle, et al., A Suite of Transgenic Driver and Reporter Mouse Lines with Enhanced Brain-Cell-Type Targeting and Functionality. Cell 174, 465–480.e22 (2018).

94. A. Mo, et al., Epigenomic Signatures of Neuronal Diversity in the Mammalian Brain. Neuron 86, 1369–1384 (2015).

95. J. Rossi, et al., Melanocortin-4 receptors expressed by cholinergic neurons regulate energy balance and glucose homeostasis. Cell Metab. 13, 195–204 (2011).

96. S. Bourane, et al., Identification of a spinal circuit for light touch and fine motor control. Cell 160, 503–515 (2015).

97. B. Duan, et al., Identification of Spinal Circuits Transmitting and Gating Mechanical Pain. Cell 159, 1417–1432 (2014).

98. K. Noben-Trauth, Q. Y. Zheng, K. R. Johnson, Association of cadherin 23 with polygenic inheritance and genetic modification of sensorineural hearing loss. Nat. Genet. 35, 21–23 (2003).

99. H. Miura, R. M. Quadros, C. B. Gurumurthy, M. Ohtsuka, Easi-CRISPR for creating knock-in and conditional knockout mouse models using long ssDNA donors. Nat. Protoc. 13, 195–215 (2018).

100. R. M. Quadros, et al., Easi-CRISPR: a robust method for one-step generation of mice carrying conditional and insertion alleles using long ssDNA donors and CRISPR ribonucleoproteins. Genome Biol. 18, 92 (2017).

101. C. S. Raymond, P. Soriano, High-efficiency FLP and PhiC31 site-specific recombination in mammalian cells. PloS One 2, e162 (2007).

102. H. Gu, Y. R. Zou, K. Rajewsky, Independent control of immunoglobulin switch recombination at individual switch regions evidenced through Cre-loxP-mediated gene targeting. Cell 73, 1155–1164 (1993).

103. F. A. Ran, et al., Genome engineering using the CRISPR-Cas9 system. Nat. Protoc. 8, 2281–2308 (2013).

104. S. Choudhary, R. Satija, Comparison and evaluation of statistical error models for scRNA-seq. Genome Biol. 23, 27 (2022).

105. C. Hafemeister, R. Satija, Normalization and variance stabilization of single-cell RNA-seq data using regularized negative binomial regression. Genome Biol. 20, 1–15 (2019).

106. T. Stuart, et al., Comprehensive Integration of Single-Cell Data. Cell 177, 1888–1902.e21 (2019).

107. J. Schindelin, et al., Fiji: an open-source platform for biological-image analysis. Nat. Methods 9, 676–682 (2012).

108. M. Linkert, et al., Metadata matters: access to image data in the real world. J. Cell Biol. 189, 777–782 (2010).

